# Arbor function of TRANSPARENT TESTA GLABRA 1 and LIGHT REGULATED WD scaffold proteins in the Arabidopsis circadian oscillators includes transcriptional repression through PSEUDO RESPONSE REGULATORS

**DOI:** 10.1101/2025.02.04.636234

**Authors:** Eva Herrero, Dora Cano-Ramirez, Beverley J. Glover, Alex A. R. Webb

**Affiliations:** Department of Plant Sciences, University of Cambridge; Current address Sainsbury Laboratory Cambridge University

## Abstract

Arabidopsis circadian oscillators contain DNA-binding proteins that function at specific times of day. In contrast, we have discovered a continuous function of WD40-repeat scaffold proteins from the TRANSPARENT TESTA GLABRA-1 (TTG1), LIGHT REGULATED WD1 and LIGHT REGULATED LWD2 subfamily (TLWD) which is essential to maintain circadian rhythms. Gene expression analyses indicate multifunctional activity of TLWD in both transcriptional activation and repression. TLWD proteins interact with an array of circadian oscillator activators and repressors that act sequentially throughout the diel cycle. While TLWD proteins were known to participate in activator complexes, our data indicate a novel role of TLWD in transcriptional repression through complex formation with repressors from the PSEUDO RESPONSE REGULATOR (PRR) family. In an analogy to mechanical clocks, TLWD scaffold proteins constitute an ever-present arbor, or spindle, to which transcription factors, which represent the cogs, bind to sustain circadian rhythms.

## Introduction

Plant circadian oscillators regulate networks of gene activity to respond appropriately to the daily rhythms of the environment, contributing to fitness(1, 2). Consequently, plant circadian oscillators control critical yield traits such as flowering time, growth, and response to environmental stress(3). Circadian oscillators contain arrays of transcription factors that are sequentially expressed through the diel cycle and orchestrate circadian rhythms of expression of over a third of the Arabidopsis transcriptome(4).

In the morning, Myb-like transcription factors contribute both to transcriptional repression, for example CIRCADIAN CLOCK ASSOCIATED (CCA1)(5) and LATE ELONGATED HYPOCOTYL (LHY)(6, 7), and to transcriptional activation, for example REVEILLE8 (RVE8)(8) in Arabidopsis. The members of the PSEUDO RESPONSE REGULATOR (PRR) family PRR9(9), PRR7(10), PRR5(11) and PRR1 (also known as TIMING OF CAB EXPRESSION1 (TOC1))(12) work sequentially as transcriptional repressors from early morning to late night(13). LUX ARRYTHMO (LUX), a member of the GOLDEN2, ARR-B, PSR1 (GARP) family, binds DNA as part of an Evening Complex of proteins to mediate transcriptional repression(14–16). Additionally, members of the TEOSINTE BRANCHED 1, CYCLOIDEA, PCF1 (TCP) family TCP20 and TCP22 contribute to the activation of *CCA1*(17, 18), whereas CCA1-HICKING EXPEDITION (CHE or TPC21) reduces *CCA1* expression(19). Overall, oscillations of expression of circadian rhythmic genes (CRGs) are orchestrated by the sequential binding of transcription factors to specific promoter *cis*-elements, resulting in either transcriptional activation or repression.

Plant circadian oscillators also feature a large array of structurally diverse scaffold proteins. Scaffold proteins do not bind directly to DNA, instead they coordinate the formation of protein complexes through their multiple binding surfaces. For instance, EARLY-FLOWERING-3 (ELF3) nucleates the Evening Complex with ELF4 and LUX(16, 20). The scaffold protein GIGANTEA (GI) promotes high amplitude oscillations of PRR5 and TOC1 by modulating the activity of the F-Box protein ZEITLUPE (ZTL)(21–23). Both ELF3 and GI function as hubs that integrate multiple environmental signals to control growth and development(14, 24, 25). In Arabidopsis, scaffold proteins of the TRANSPARENT-TESTA-GLABRA-1 (TTG1) and LIGHT-REGULATED-WD40-repeat (LWD) family (TLWD) are an essential part of the circadian oscillator besides controlling many other aspects of plant biology(26–28). Functional divergence in the TLWD family arose at the base of the seed plant lineage(27). In Arabidopsis, TTG1 itself is an integral part of the Myb-bHLH-WDR (MBW) complexes that control pigment biosynthesis and epidermal cell differentiation(28) while LWD1 and LWD2 control circadian period and light entrainment(26). Importantly, LWD1, LWD2 and TTG1 all redundantly contribute to sustain circadian rhythms(27) since the *ttg1 lwd1 lwd2* triple mutant has arrhythmic circadian outputs such as gene expression and leaf movement. LWDs are required for TCP-mediated activation of *CCA1* expression(17, 18). However, loss of TCP function does not lead to arrhythmia but only to changes in amplitude of *CCA1* expression, suggesting additional roles of TLWD in circadian oscillators(17, 18).

Here we report that TLWD proteins have a unique multifunctional role in the circadian oscillator. We show that both LWDs and TTG1 interact with transcriptional repressors of the Myb-like and PRR families, and with transcriptional activators of the TCP family. Specifically, we find that transcriptional repression mediated by TLWD-PRR complexes is essential to sustain circadian oscillations and regulate circadian outputs. TLWD proteins represent a new category of circadian oscillator component, which, unlike most plant circadian oscillator proteins, are not temporally restricted in abundance. Because of their continuous presence, and their ability to bind multiple different transcriptional repressors and activators throughout the 24 h cycle, we describe TLWD proteins as arbor proteins, an arbor being the spindle of a mechanical clock or watch to which the cogs and gears are attached.

## Results

### TLWD function is required for both activation and repression of circadian rhythmic genes

We previously reported that TLWD function is essential to sustain circadian oscillations of *CCA1* gene expression and leaf movements in Arabidopsis(27). To understand more widely the impact of LWDs and TTG1 in transcriptional regulation, we measured transcript abundance by RNAseq under diel cycles (12 hr light / 12 hr dark), sampling 10-day-old seedlings in the middle of the day (ZT6) and in the middle of the night (ZT18) to capture temporal regulation. We measured differences in transcript abundance between wildtype and *ttg1*, *lwd1 lwd2* double and *ttg1 lwd1 lwd2* triple mutants. We considered differentially expressed genes (DEG) as those with a Log2^FC^ >1.5 between the mutant and the wild type at the fixed time point. Only a small number of genes were downregulated in the *ttg1* mutant compared to wild type at ZT6. These corresponded to known direct targets of MBW complexes involved in flavonoid biosynthesis(29) (Fig. S1A, Table S1), demonstrating the ability of our approach to identify direct targets of TLWD regulation of transcriptional activity. These direct targets of MBW complexes were also downregulated in the *ttg1 lwd1 lwd2* triple mutant but not in the *lwd1 lwd2* double mutant (Fig. S1A, Table S1), consistent with LWDs not complementing the TTG1 functions mediated by MBW complexes(27). The majority of DEGs in the *lwd1 lwd2* and *ttg1 lwd1 lwd2* mutants were known circadian rhythmic genes (CRG)^4^, indicating that the predominant activity of TLWD proteins in young seedlings is circadian regulation (*lwd1 lwd2,* Day: 97% of the down-regulated and 80% of the upregulated genes, Night: 97% of the upregulated genes and only 1 gene downregulated; *ttg1 lwd1 lwd2,* Day: 57% of the downregulated genes and 75% of the upregulated genes, Night: 45% of the downregulated genes and 79% of the upregulated gene, Figure 1A, Table S1). Consistent with the lack of circadian phenotypes in the single *ttg1*(27) mutant, we did not find a defect in circadian regulated gene expression in *ttg1* (Table S1).

**Figure 1.**
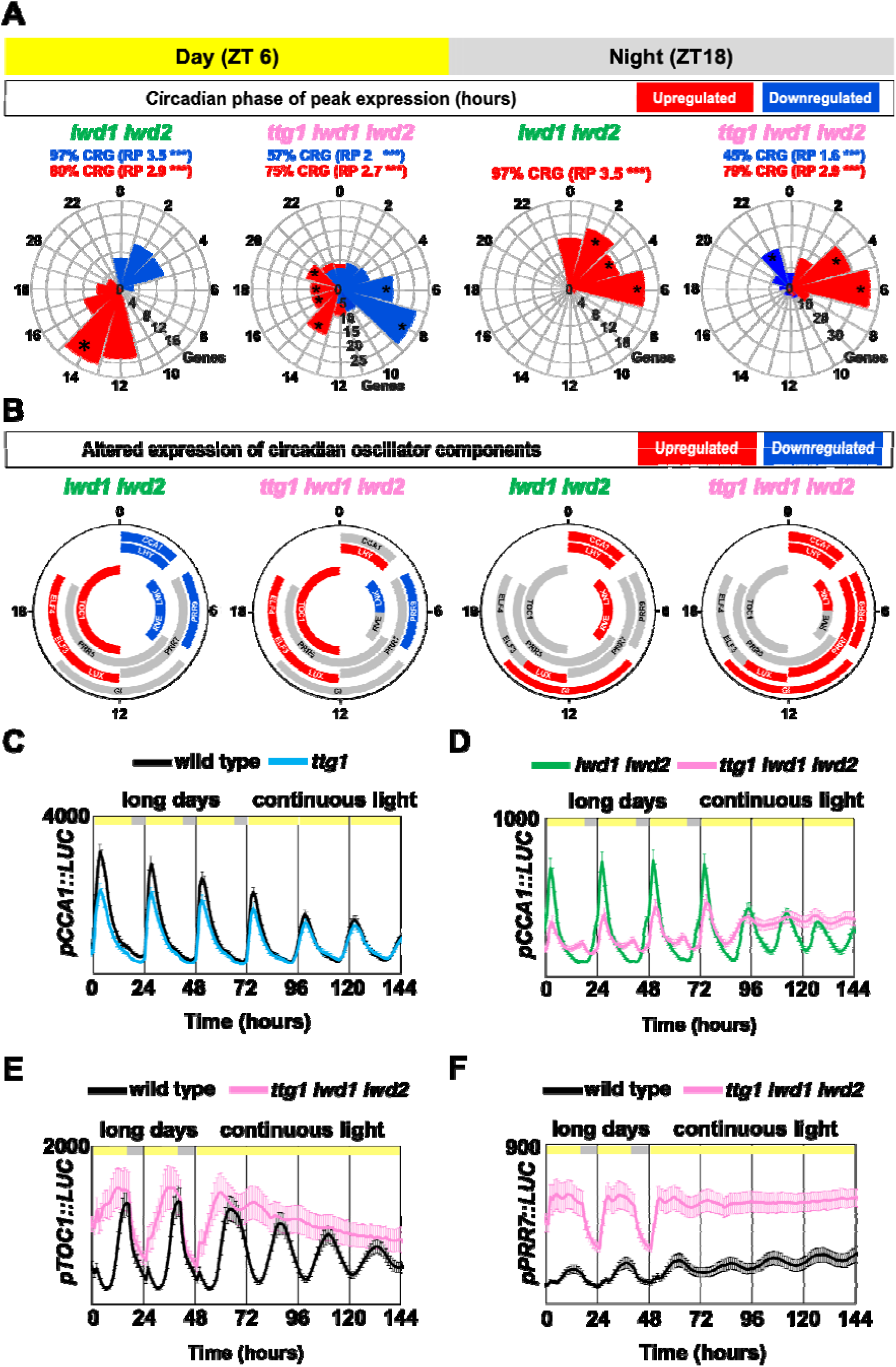
TLWD proteins regulate circadian rhythms of gene expression. (A) Percentage of genes upregulated (red) or downregulated (blue) that are circadian rhythmic (CRG) in *lwd1 lwd2* (green) and *ttg1 lwd1 lwd2* (pink) mutants during day (ZT6) or night (ZT18) and distribution of their usual circadian phase of expression in wild type. RP: representation factor. Asterisks depict statistically significant enrichment of the circadian phase. Figure data in Table S1. (B) Circadian oscillator components upregulated (red) or downregulated (blue) relative to wild type in *lwd1 lwd2* (green) and *ttg1 lwd1 lwd2* (pink) mutants during day (ZT6) or night (ZT18). Oscillator genes are depicted according to their usual expression pattern through the diel cycle. Genes with no change in expression are depicted in grey. See Fig. S1C for normalized expression levels. (C-D) Oscillations of *pCCA1::LUC* in wild type and *ttg1* mutants (C) and *lwd1 lwd2* and *ttg1 lwd1 lwd2* mutants under diel cycles (16h light / 8h dark) and continuous light (free running) (D). Y axis depicts mean bioluminescence (counts in 800 sec) of cluster of seedlings, n=24 per genotype. Note the different maximum Y-axis values. (E-F) Oscillations of *pTOC1::LUC* (E) and *pPRR7::LUC* (F) in wild type and *ttg1 lwd1 lwd2* mutants under diel cycles (16h light / 8h dark) and continuous light (free running). Y axis depicts mean bioluminescence (counts in 800 sec), of *pTOC1::LUC* (wild type n=8, *ttg1 lwd1 lwd2* n=8) and *pPRR7::LUC* (wild type n=8, *ttg1 lwd1 lwd2* n=12). Error bars in C-F depict SEM.

The trend of misregulation of CRGs was comparable between *lwd1 lwd2* and *ttg1 lwd1 lwd2,* but was more pronounced in the triple mutant, confirming genetic redundancy between TLWD proteins (Fig.S1B). In the day, at ZT6, from the 97 CRG downregulated in *ttg1 lwd1 lwd2* mutants (Log2^FC^ >-1.5), 62 were also downregulated in *lwd1 lwd2* (Log2^FC^ >-0.5) (Table S1). We used the CAST-R database(30) to analyse the wild type circadian phase of expression of misregulated CRGs in *lwd1 lwd2* and *ttg1 lwd1 lwd2* mutants (Figure 1A, Table S1). Downregulated CRGs at ZT6 in *lwd1 lwd2* and *ttg1 lwd1 lwd2* usually had a morning circadian phase (ZT0-6). In addition, 47 out of 97 CRGs downregulated in *ttg1 lwd1 lwd2* usually had a late morning circadian phase ZT6-ZT8 (Fold enrichment 3.89-3.78, p-value <0.05). The lack of activation of CRGs in the morning in *TLWD* mutants is consistent with the known role of LWDs in activating gene expression in the morning(17, 18)

Whilst the role of LWDS in activation of gene expression is known, we also discovered an important but previously overlooked role of TLWD in transcriptional repression at different times of the day. We found defects of circadian-associated transcriptional repression both in the day and in the night (Figure 1A, Table S1). The majority of CRGs upregulated in *ttg1 lwd1 lwd2 (*Log2^FC^ >1.5) were also upregulated in *lwd1 lwd2 (*Log2^FC^ >0.5), both at ZT6 (66/86) and at ZT18 (87/111) (Table S1). CRGs upregulated in *lwd1 lwd2* at ZT6 usually had a circadian phase of expression of ZT14 (Figure 1A, Table S1; Fold enrichment ZT6=1.92, p value < 0.005). CRGs upregulated in *ttg1 lwd1 lwd2* at ZT6 have a usual circadian phase of expression of ZT14-ZT20 (Figure 1A, Table S1; ZT (Fold enrichment): ZT14 (2.4), ZT16 (2.4), ZT18 (2), ZT20 (1.9), p-value < 0.05). Conversely, CRGs upregulated in *lwd1 lwd2* and *ttg1 lwd1 lwd2* mutants at ZT18 normally have a morning circadian phase of expression (ZT2 to ZT6) (Figure 1A, Table S1, ZT (Fold enrichment): *lwd1 lwd2* ZT2 (1.8), ZT4 (2.8), ZT6 (4.6); *ttg1 lwd1 lwd2* ZT4 (4.4), ZT6 (6.3), p value < 0.05).Most components of the Arabidopsis circadian oscillator were misexpressed in *lwd1 lwd2* and *ttg1 lwd1 lwd2* mutants. Morning genes *CCA1*, *PRR9* and *LNK2* were downregulated at ZT6 and upregulated at ZT18 in comparison to wild type, whereas evening expressed genes *TOC1* and *ELF4* were upregulated at ZT6. *LUX* was upregulated both at ZT6 and ZT18 (Figure 1B, Fig. S1C and Table S1). Taken together, we have identified a dual role of TLWD in the circadian oscillator mediating transcriptional activation in the morning and transcriptional repression in the morning and at night. To look more closely at the temporal aspects of the effects of TLWD function we analysed rhythms of activity of *CCA1*, *TOC1* and *PRR7* promoters fused to firefly luciferase (LUC) both under diel and free-running conditions. We previously reported that the *ttg1* mutant under free running conditions does not have a defect in *pCCA1::LUC* activity (27). In our current analysis, we also did not see a defect under diel cycles either (Figure 1C), confirming that TTG1 function is redundant to that of the LWDs^27^. Conversely, both *lwd1 lwd2* and *ttg1 lwd1 lwd2* mutants had reduced amplitude of *pCCA1::LUC* activity under diel cycles (Figure 1D). In *ttg1 lwd1 lwd2* there was a second peak of *pCCA1::LUC* in the second half of the day. This second peak was only apparent under long-day photoperiods (LD) (Figure 1D) but not under neutral or short-day photoperiods (SD) (Fig. S2), indicating a lack of repression in the second half of the diel cycle which allows light to induce *CCA1* promoter activity. As previously shown (27), under free-running conditions, oscillations of *pCCA1::LUC* became arrhythmic. Similarly to the lack of repression under diel LD, *pCCA1::LUC* expression increased from subjective dusk and remained high in the *ttg1 lwd1 lwd2* mutant (Figure 1D). These data indicate that under diel cycles TLWD have a dual function in sustaining *CCA1* oscillations. In the morning LWDs activate *CCA1* expression (17, 18) and in the second part of the day both LWDs and TTG1 repress *CCA1* expression. These data uncover a lack of repression of *pCCA1::LUC* in the *ttg1 lwd1 lwd2* mutant at dusk under LD. We then tested the expression of other clock-regulated promoters and found a role for repression at other points in the daily cycle. (Figure 1E-F). In wildtype plants the activity of *pPRR7* and *pTOC1* peaked at ZT12 and ZT16 (dusk) respectively, consistent with previous reports. However, in *ttg1 lwd1 lwd2* plants, the activity of both *pPRR7* and *pTOC1* started to increase at dawn and remained high until dusk indicating a lack of repression in the morning. Similarly to *pCCA1*, under continuous light, the expression of both *pPRR7* and *pTOC1* remained high and became arrhythmic (Figure 1E-F). Taken together, our analysis of circadian regulated gene expression and promoter activity under diel cycles discovers an essential role of TLWD in circadian mediated transcriptional repression throughout the diel cycle.

### TLWD proteins are present through the diel cycle

The abundance and/or activity of plant circadian oscillator components is restricted to specific times of the day. However, our data are consistent with a continuous function of TLWD proteins through the diel cycle. To test whether LWD1 and TTG1 proteins are present through the diel cycle, we generated complementation lines of *lwd1 lwd2* and *ttg1* mutants with LWD1-Venus or Venus-TTG1 flanked by their respective promoters and downstream sequence. For LWD1, we selected complementation lines that fully complemented *lwd1 lwd2 pCCA1::LUC* oscillations (Fig. S3). For TTG1, we selected complementation lines that restored trichomes and seed coat pigmentation of *ttg1* mutants (Fig. S4), indicating that the addition of the Venus tag did not compromise protein function. We confirmed the nuclear localization of LWD1-Venus and TTG1-Venus in the corresponding complementation lines (Fig. S3-S4). To confirm that our approach can detect oscillations in protein abundance, we included a previously described *pCCA1::CCA1-YFP*(*31*) line in our analysis. We sampled 10-day-old seedlings grown under diel cycles (12h light / 12h dark) every 4 hours during one diel cycle and prepared whole protein extracts. We used the same anti-GFP antibody for detecting CCA1, LWD1 and TTG1 by western blot. CCA1-YFP protein abundance was restricted to morning (ZT0-ZT8) peaking at ZT4 (Figure 2A). In contrast, both LWD1-Venus and TTG1-Venus were present throughout the day and night without a clear reproducible oscillatory pattern (Figure 2 B-C). To increase the temporal resolution of LWD1 and TTG1 detection we generated LWD1-LUC and TTG1-LUC complementation lines in the *lwd1 lwd2* and *ttg1* mutant lines driven by the corresponding endogenous promoter (see complementation data in Fig. S4C and western blot under diel cycles Fig. S5). We detected low amplitude oscillations of luminescence in both LWD1-LUC and TTG1-LUC with a first peak at dawn and a second peak at dusk under diel cycles, and protein levels decreasing during the night (Figure 2D-E). Low amplitude oscillations persisted under continuous light for both proteins. The fold change of LWD1 and TTG1 abundance was comparable in the western blots (LWD1 ∼ 2.1 and TTG1 ∼ 1.7) and LUC activity (LWD1 ∼ 1.8 and TTG1 ∼ 1.6) (Figure 2 B-E), which is two orders of magnitude lower than oscillation of CCA1-YFP abundance (∼ 280) Figure 2A). Taken together, both LWD1 and TTG1 proteins are present through the diel cycle (unlike CCA1) and have extremely low amplitude oscillations of abundance peaking at subjective dusk under free-running conditions.

**Figure 2.**
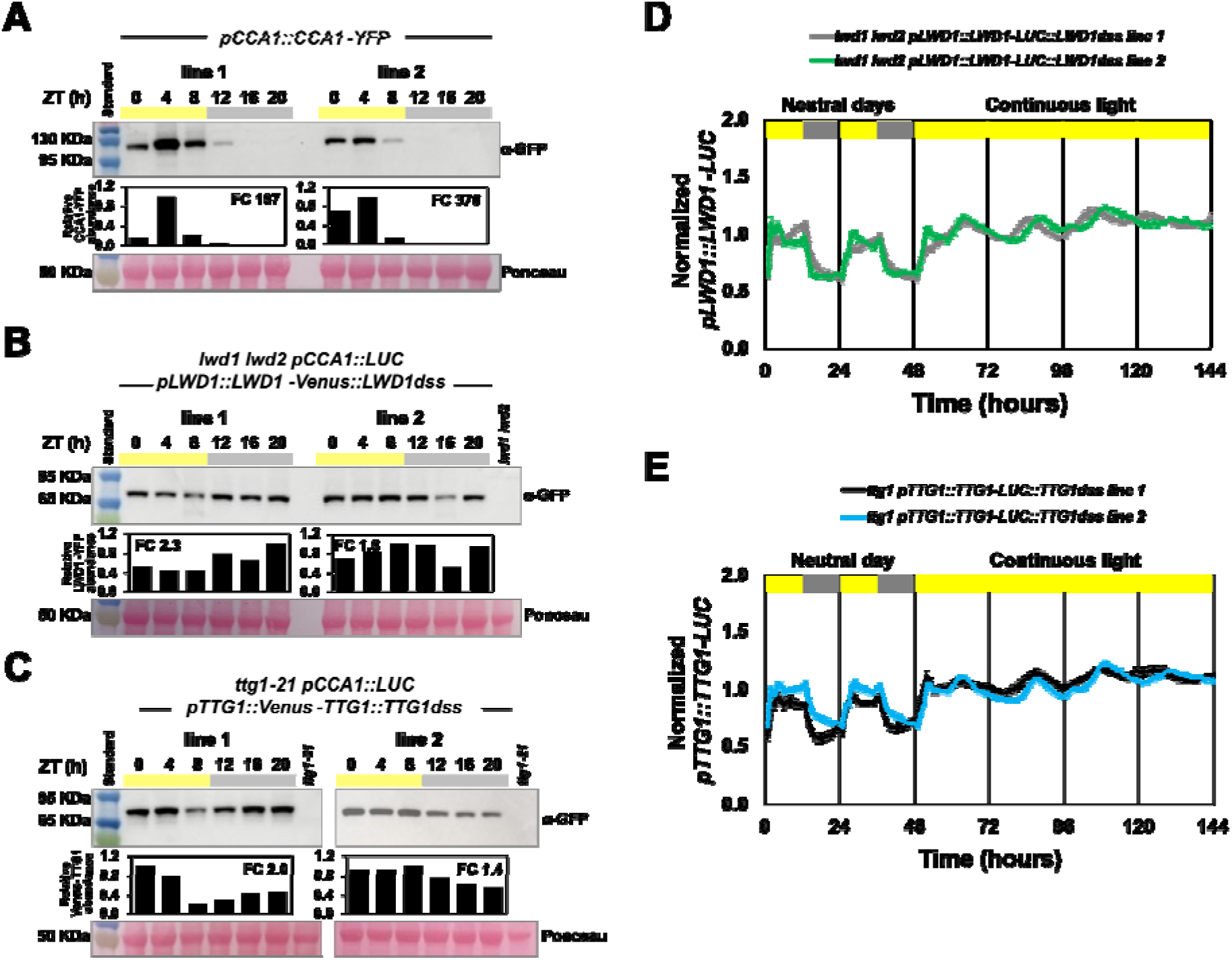
TLWD proteins are present throughout the diel cycle. Representative western blots from protein extracts of (A) *pCCA1::CCA1-YFP* (B) *lwd1 lwd2 pCCA1::LUC pLWD1::LWD1-Venus* and (C) *ttg1 pCCA1::LUC pTTG1::Venus-TTG1*. Seedlings from two independent lines (Line 1 and Line 2) were grown for 10 days under diel cycles (12h light / 12h dark) and harvested at the indicated time points. Expected molecular weight: CCA1-YFP (95 KDa), LWD1-Venus (67 KDa) and Venus-TTG1 (67 KDa). Bar graphs show protein abundance relative to maximum. Ponceau staining shown as loading control, 10 µg of protein extract was loaded for each timepoint. (D) and (E) Rhythms of *pLWD1::LWD1-LUC* and *pTTG1::TTG1-LUC* under diel cycles (12h light /12h dark) and continuous light. Error bars depict SEM. *lwd1 lwd2 pLWD1::LWD1-LUC* line 1(n=16) and 2 (n=21). *ttg1 pTTG1::TTG1-LUC* line 1 (n=14) and 2 (n=24).

### TLWD proteins interact with both circadian oscillator activators and repressors

*ttg1 lwd1 lwd2* mutants are arrhythmic(27). However, why this might be the case has been unclear because the only known interactors with the TLWD proteins were the TCP proteins. The effects of *ttg1 lwd1 lwd2* cannot be solely due to lack of transcriptional activity of LWD-TCP complexes at dawn, since *tcp* mutants only have defects in amplitude of *pCCA1* activity and retain full rhythmicity(17). Our transcriptome and promoter activity data indicated other potential roles for TLWD proteins in transcriptional repression (Figure 1), which we hypothesized might explain why the triple mutant is arrhythmic. We tested whether TLWD proteins interact with additional clock transcription factors, particularly the transcriptional repressors that are active throughout the day. In yeast 2 hybrid assays, TCP20, TCP22 and CHE interacted with LWD1 and LWD2, as previously reported(17). We also found interaction of TCP20, TCP22 and CHE with TTG1 (Fig. S6). Additionally, LWD1, LWD2 and TTG1 all interacted with PRR5 (Figure 3A). TTG1 itself also interacted with PRR9, PRR7, TOC1 and PRR3 (Figure 3A). Finally, TTG1 interacted with Myb-like transcriptional repressors CCA1, LHY and RVE1, but not with the activator RVE8 (Fig. S7A). These data are consistent with a continuous function of TLWD proteins throughout the circadian cycle, forming complexes with different transcriptional repressors and activators that act sequentially through the day and night in a sequence that starts with CCA1/LHY after dawn, moves through the PRRs and ends with the TCPs just around dawn. To test predictions of the yeast two hybrid data *in planta* we performed co-immunoprecipitation in *N. benthamiana* using transient expression. Both LWD1 and TTG1 interacted with PRR9, PRR7 and PRR5. Also, LWD1 interacted with TOC1 (Figure 3B-C). Differences in the interaction ability between LWD1 and TTG1 with PRR proteins in our Y2H or Co-IP experiments may be caused by differences in posttranslational modifications between both systems. In summary, we found that both LWD1 and TTG1 interact with multiple PRR proteins *in planta,* implying that multiple different TLWD-PRR complexes may be formed during progress through the diel cycle.

**Figure 3.**
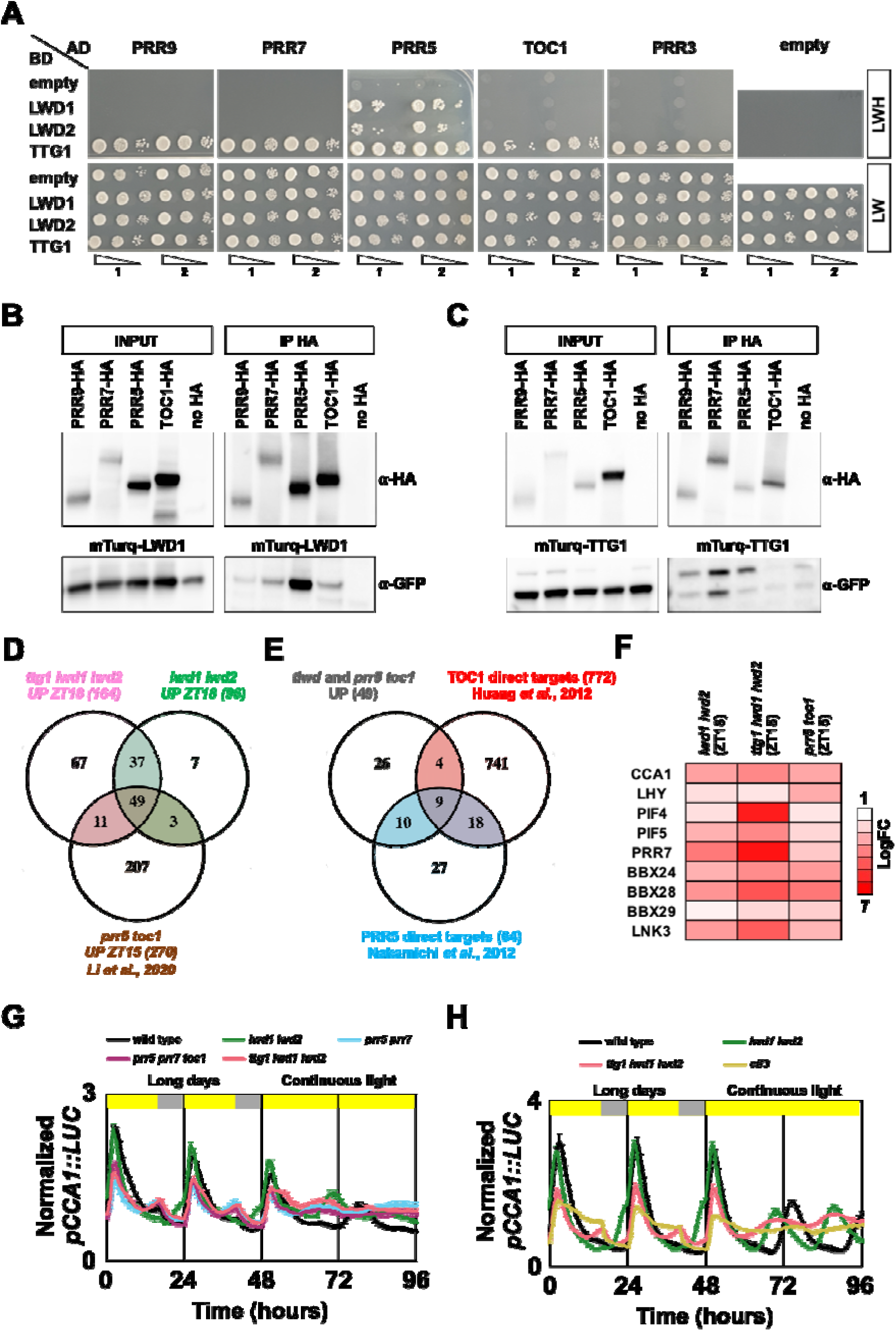
TLWD proteins form complexes with PRRs. (A) Y2H protein interaction assay: 3 serial dilutions (1:10) of two representative colonies. Growth in LHW media indicates interaction between protein pair. (B) and (C) Co-immunoprecipitation of LWD1 (B) and TTG1 (C) with PRR proteins from *N. benthamiana* extracts. (D) Venn diagram depicts overlap of CRGs upregulated relative to wildtype in *lwd1 lwd2* (ZT18), *ttg1 lwd1 lwd2* (ZT18), and *prr5 toc1 (ZT15)^32^. (E)* Venn diagram depicts overlap between the 49 CRGs upregulated in both *tlwd* and *prr* mutants and the direct targets of TOC1 and PRR5 as established by ChIPseq^12,33^. (F) Fold change of expression of potential TLWD-PRR complex targets in corresponding transcriptome analysis. (G-H) Comparison of *pCCA1::LUC* oscillations in (G) *tlwd* and *prr* mutants or (H) *tlwd* and *elf3* mutants. Y axis depicts mean normalized bioluminescence of clusters of seedlings. Error bars depict SEM, genotype (n): in (G) wild type (15), *lwd1 lwd2* (12), *prr5 prr7* (16), *prr5 prr7 toc1* (16), *ttg1 lwd1 lwd2* (16) and in (H) wild type (20), *lwd1 lwd2* (11), *ttg1 lwd1 lwd2* (23), *elf3* (20).

Similarly to *ttg1 lwd1 lwd2* mutants, triple *prr* loss of function mutants are arrhythmic(32). Hence, it is possible that loss of PRR repression, in addition to the previously identified loss of TCP activation(17), might account for the arrhythmicity in *ttg1 lwd1 lwd2* triple loss of function mutants. We reasoned that potential direct targets of TLWD-PRR complexes would be upregulated both in higher order *tlwd* and *prr* mutants due to a common defect in transcriptional repression. Hence, we compared CRGs that were upregulated both in our RNAseq experiment at ZT18 in both *lwd1 lwd2* and *ttg1 lwd1 lwd2* with CRGs upregulated in *prr5 toc1* mutants at a similar time ZT15 (Log2^FC^>1)(33). We found that 49 genes were shared between these two datasets (Fold enrichment (p value): *lwd1 lwd2* 62.07 (p 5.9 e^-63^) and *ttg1 lwd1 lwd2* 36.33 (5.9 e^-63^) (Figure 3D). Of these 49 genes, 9 were also direct targets of both PRR5(34) and TOC1(12) repression as established by previous CHIP-seq experiments (Figure 3E, F). Both *PRR7* and *CCA1* were potential targets of TLWD-PRR complexes (Figure 3F). Consistently, we found a lack of repression of both *pCCA1* and *pPRR7* in *ttg1 lwd1 lwd2* mutants (Figure 1D and 1F). Notably, we found that under LD both *prr7 prr5* and *prr7 prr5 toc1* had a second peak of *pCCA1* activity in the second half of the diel cycle and under free-running conditions *pCCA1* activity remained high (Figure 3G), phenocopying the characteristic *pCCA1* activity of *ttg1 lwd1 lwd2* mutants (Figure 1D). The arrhythmic mutant *elf3* did not phenocopy *ttg1 lwd1 lwd2* aberrant *pCCA1* oscillations under diel cycles (Figure 3H), supporting the idea that loss of repressive activity of PRR complexes leads to the second peak of *pCCA1* activity in *ttg1 lwd1 lwd2* triple mutant plants.

The Y2H data also suggested that TLWD proteins interact with Myb-like transcription factors (Fig. S7A). To test whether a defect in CCA1/LHY activity might contribute to the arrhythmicity of *ttg1 lwd1 lwd2* mutants, in addition to our analysis of *pCCA1* (Figure1C, D), we compared rhythms of *pTOC1::LUC* under diel cycles and free running conditions in *ttg1 lwd1 lwd2* and *lhy cca1* mutants as the *TOC1* promoter is a direct target of CCA1/LHY repression(35). We also included in the comparison the *elf3-1* mutant which is arrhythmic and has low levels of *CCA1/LHY* expression(20, 36), similar to *ttg1 lwd1 lwd2* mutants(27) (Figure 1D). Wild type plants had a small transient peak of *pTOC1* at dawn and a second main peak around dusk. In contrast, *ttg1 lwd1 lwd2* and *lhy cca1* mutants all had high levels of *pTOC1* expression after the dawn peak until dusk where *pTOC1* levels decreased (Fig S7B). These data are consistent with TLWD contributing to CCA1/LHY repression of *TOC1*. The *elf3-1* mutant was different since *pTOC1* was clearly repressed after the dawn peak and the second peak was only slightly advanced (Fig. S7B).

### TLWD proteins control growth through repression of *PIF4/5* expression

Arabidopsis circadian oscillators provide a temporal framework to alter growth in response to environmental signals by generating circadian rhythms of expression of the growth promoting transcription factors *PHYTOCHROME INTERACTING FACTOR4* (*PIF4*) and *PIF5* (16, 37). We have identified both *PIF4* and *PIF5* as potential targets of TLWD-PRR complexes (Figure 3F). In our transcriptome analysis, we found that *PIF4* and *PIF5* are upregulated specifically in the *ttg1 lwd1 lwd2* mutant at night (ZT18), together with various direct targets of PIF4 and PIF5 activation such as *CYCLING DOF FACTOR 5 (CDF5)* and *ARABIDOPSIS THALIANA HOMEOBOX PROTEIN 2 (ATBH2)* (Figure 4A, Fig. S8A and Table S1). Since a role of TLWD in controlling growth has not been previously described, we investigated hypocotyl length of TLWD mutants under both SD and LD. Consistent with the expression of *PIF4/5* and target genes (Figure 4A, Fig. S8A), we found that *ttg1 lwd1 lwd2* mutants have longer hypocotyls than wildtype both under SD and LD (Figure 4B, Fig. S8B-C). In contrast, hypocotyl length in *lwd1 lwd2* double mutants and *ttg1* single mutants was like wild type, indicating that all three TLWD proteins work redundantly to repress hypocotyl growth. We found that *ttg1 lwd1 lwd2* had shorter hypocotyls in comparison with the other circadian arrhythmic mutants *prr7 prr5 toc1, prr9 prr7 prr5* and *elf3* (Figure 4B, Fig. S8B-C), suggesting additional effects of the loss of PRR or ELF3 function besides circadian arrhythmicity. To test whether expression of *PIF4*, *PIF5* and *CDF5* correlated with hypocotyl length, we compared the expression of *tlwd* mutants and *prr7 prr5 toc1* mutants under SD. (Figure 4C). Consistent with a role of TLWD in transcriptional repression at night, *PIF4*, *PIF5* and *CDF5* were upregulated in *ttg1 lwd1 lwd2* plants particularly in the second half of the diel cycle (ZT12-21). Expression levels were lower in *ttg1 lwd2 lw2* mutants than in *prr7 prr5 toc1* plants, which correlated with their respective hypocotyl lengths (Figure 4C). In the *lwd1 lwd2* mutant, *PIF4* levels were similar to wild type but *PIF5* expression was increased at the end of the night consistent with our transcriptome analysis (Figure 4A). Beyond promoting hypocotyl elongation, an increase in *PIF4* expression causes elongated petioles and smaller leaves when grown under SD(38). We found that *ttg1 lwd1 lwd2* plants had elongated petioles and small leaves similar to *PIF4-OX* and *elf3* mutants, whereas both *ttg1* and *lwd1 lwd2* plants were similar to wild type (Fig. S8C). Taken together, these data show that TLWD controls growth by contributing to the repression of PIF4/5 to control growth.

**Figure 4.**
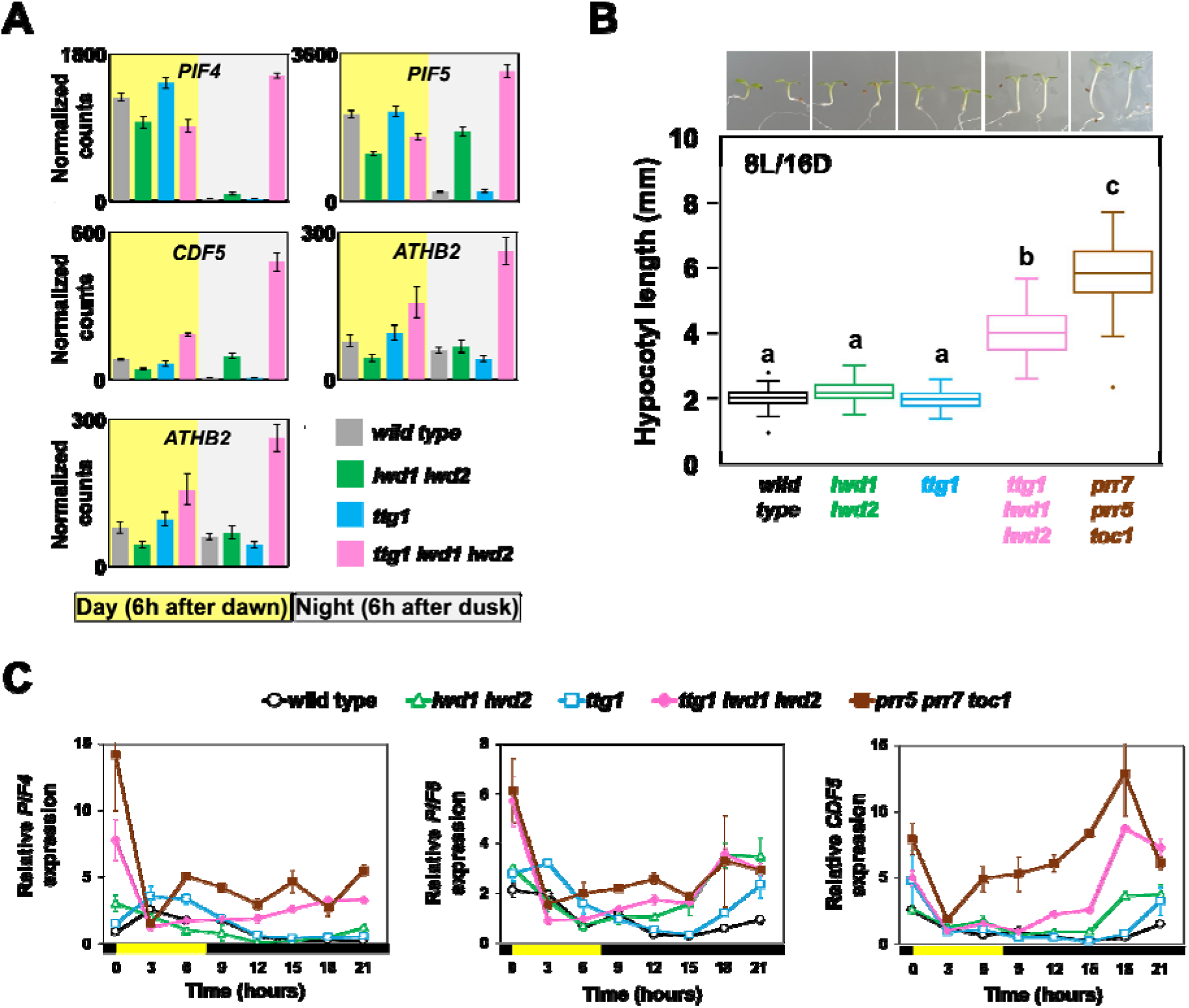
TLWD control growth by repressing *PIF4* and *PIF5* expression at night. (A) Normalized expression of *PIF4*, *PIF5* and their gene targets in wildtype, *lwd1 lwd2*, *ttg1* and *ttg1 lwd1 lwd2* mutants as measured by RNAseq. Normalized counts were obtained with Deseq2 (Table S1). Error bars depict SEM, n=3. (B) Representative seedlings and mean hypocotyl length of plants grown for 7 days in SD (8h light / 16h dark) of white light. Error bars depict SEM; n=wild type (54), *lwd1 lwd2* (47), *ttg1* (55), *ttg1 lwd1 lwd2* (46) and *prr5 prr7 toc1* (58). Letters depict genotypes with statistically different average length with the Tukey Kramer test (D) Expression profile of *PIF4, PIF5* and *CDF5* under SD. Expression values are normalized to both *PP2A* and *UBQ10* expression and plotted relative to average wild type expression through the diel cycle. Error bars depict SEM, n=3.

## Discussion

### Continuous function of TLWD in Arabidopsis circadian oscillators

Our work highlights the importance of scaffold proteins in circadian oscillators. In the context of transcriptional regulation, scaffold proteins mediate the formation of multiprotein complexes which are essential for the function of transcription factors(16, 20). In Arabidopsis, the window of activity of the scaffold protein GI expands between ZT8-ZT12 under SD and ZT4-16 under LD (39, 40) when it forms complexes with ZTL and modulates TOC1 abundance(23, 40, 41). On the other hand, the ELF3 scaffold protein nucleates the evening complex later in the diel cycle, between ZT10 and ZT20(16, 42) depending on photoperiod. In contrast to the time-restricted activity of GI or ELF3 scaffolds, we have found a unique continuous activity of TLWD proteins in the circadian oscillator by forming complexes with multiple transcription factors of different families (Figure 3, Fig. S6-S7) that work sequentially to control either the activation or repression of target genes (Figure 5). Consistent with a continuous function of TLWD, we previously showed that continuous expression of *LWD1* does not alter circadian rhythms of *pCCA1* activity(27). Therefore, the arrhythmic phenotype of *ttg1 lwd1 lwd2* mutants is a composite of sequential loss of transcriptional repression and activation activities of the binding partners of TLWD proteins throughout the diel cycle, including TCPs, CCA1/LHY, PRR5,7 and 9 and TOC1. TLWD proteins are always detectable and have very low amplitude oscillations with maximum abundance around dusk (Figure 2). In contrast, a recent study estimated that the protein levels of the circadian clock oscillator transcription factors CCA1, LHY, PRR7 and TOC1 oscillate with an order of magnitude larger amplitude ranging from 12 to 84-fold(43). On the contrary, in the same study ELF3 was found to have low amplitude oscillations in protein abundance with an estimated fold change of 1.7(43), similar to TTG1 and LWD1 (Figure 2). The combination of high amplitude oscillations of transcription factor abundance with low amplitude oscillation of scaffolds might contribute to the robustness of circadian rhythms of gene expression by ensuring appropriate transcription factor-scaffold complexes are sequentially assembled through the day. Our data suggest TLWD might sequentially bind oscillator repressors and activities throughout the entire cycle, unlike the proposed temporally restricted activities of ELF3 and GI.

**Figure 5.**
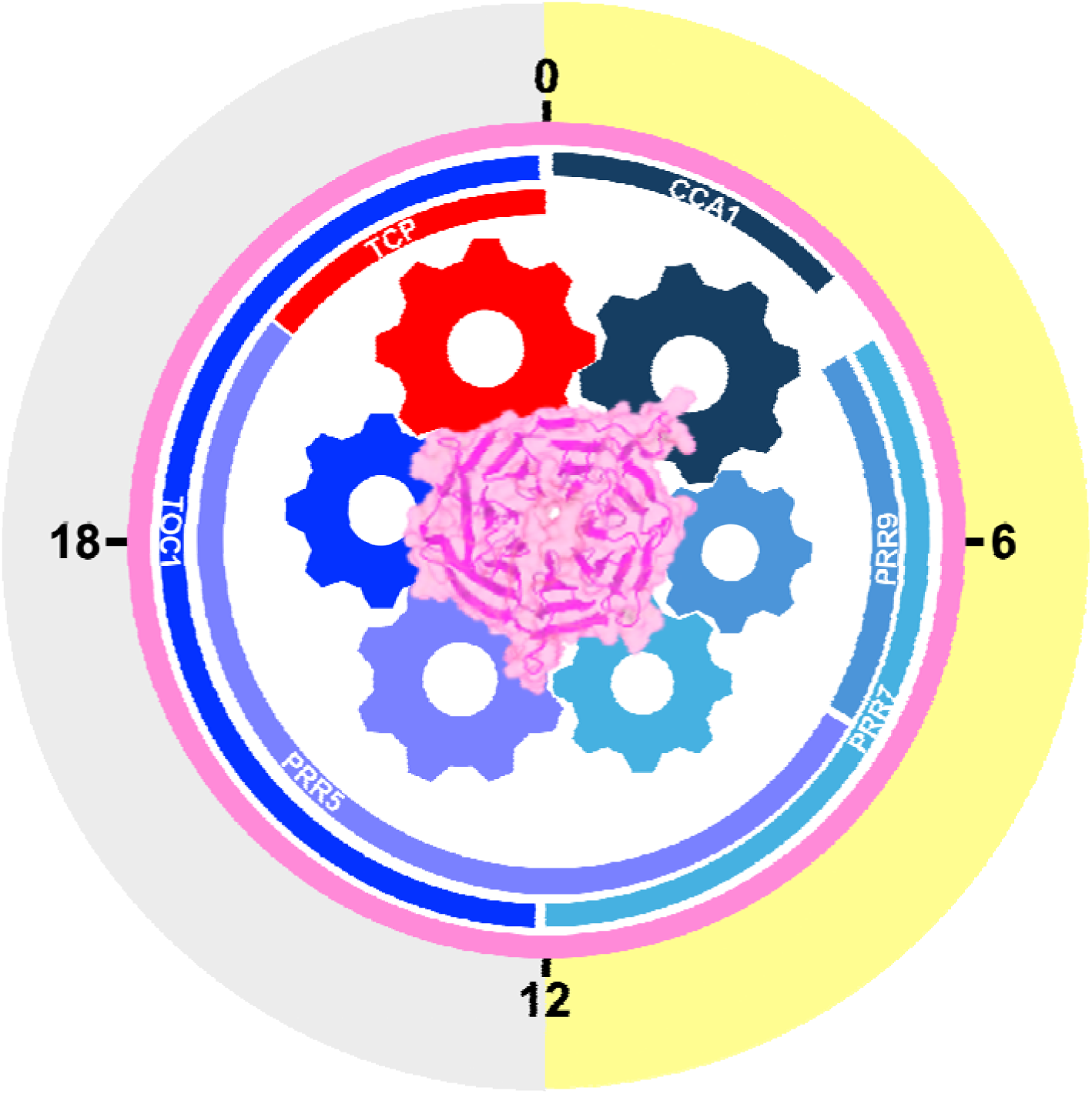
TLWD are arbor proteins in the Arabidopsis circadian oscillator. TLWD provides a continuous platform for the sequential formation of activator and repressor complexes in the Arabidopsis circadian oscillator. The model depicts sequential temporal complex formation between TLWD (central arbor) and clock transcription factors (outer wheels). TLWD structure was generated by Alphafold and illustrated with ChimeraX. Interaction surfaces between TLWD and TF are not based on structural information. Outer bars depict timing of activity of TLWD interacting clock transcription factors (red=activators, blue=repressors, pink=TLWD).

### Novel function of TLWD proteins in circadian oscillator transcriptional repression

Analysing circadian gene expression under diel cycles allowed us to compare circadian arrhythmic mutants and identify specific activation and repression defects. We found that in *ttg1 lwd1 lwd2* mutants lack of repression correlated with Myb-like transcription factor repression during the day and with PRR-mediated repression at night (Figure 1,4 and Fig. S7). Our data are consistent with the Evening Complex not being compromised by the absence of *tlwd* function because *elf3* mutants have different diel oscillations of both *pCCA1* and *pTOC1* to *ttg1 lwd1 lwd2* mutants (Figure 4 and Fig. S7).

Until now, the activity of TTG1 and LWDs has been associated with transcriptional activation, through MBW complexes in the case of TTG1 and through LWD-TCP complexes(17, 18, 28, 44). However, our data indicate that it is the lack of repression that causes arrhythmicity in the absence of TLWD function. Blue light dependent LWD1-TCP-CRY2 association with the *CCA1* promoter at dawn mediates *CCA1* transcriptional activation(17, 18). Notably, in the *tcp20/tcp22* double mutant, LWD1 is still associated with the *CCA1* promoter, indicating that additional transcription factors mediate its binding. Our data support a direct role of TLWD-PRR complexes in repressing *CCA1,* which is most apparent in the second half of the diel cycle. High order *prr* mutants phenocopy the aberrant expression of *pCCA1::LUC* in *ttg1 lwd1 lwd2* mutants (Figure 1D and 4G), including the second *pCCA1::LUC* peak under LD.

We identified a function of TLWD in controlling hypocotyl growth by repressing *PIF4* and *PIF5* expression, possibly via TLWD-PRR complexes (Figure 4F and 5, Fig. S8). Notably, *ttg1 lwd1 lwd2* mutants have lower levels of *PIF4* expression and shorter hypocotyls when compared to *prr5 prr7 toc1* mutants (Figure 4). In the *tlwd* null mutant, lower levels of *PIF4* expression may be caused by increased repression by the evening complex since we found upregulation of *ELF4*, *ELF3* and *LUX* in the morning (ZT6, Fig. S1C).

CCA1/LHY directly repress the expression of *PRR* genes(45). We found that *lhy cca1* and *ttg1 lwd1 lwd2* have a similar de-repression of *pTOC1* rhythms during the day (Figure 1, Fig. S7). Notably, Wang *et al.* previously found that LWD1 associates with the *TOC1* promoter at ZT0 and at ZT6, when *TOC1* is repressed, but not at ZT12, when *TOC1* expression is highest(26). This supports a direct role of TLWD proteins in the repression of *TOC1* via TLWD-CCA1 complexes. In addition, LWD1 binds to the *PRR9* promoter at ZT0, 6 and 12, which coincides with both high and low levels of *PRR9* expression(26). We found that *PRR9* was downregulated at ZT6 in both *lwd1 lwd2* and *ttg1 lwd1 lwd2* mutants (Figure 1B, Fig. S1), further supporting a continuous function of TLWD proteins which involves both activation and repression of circadian regulated genes.

### TLWD complexes as a model for dynamic transcriptional regulation

How TLWD proteins contribute to the function of circadian oscillator transcription factors is still poorly understood. We propose that TLWD scaffolds might provide a continuous platform that connects the activity of different transcription factor families that bind sequentially to the same circadian-regulated promoters such as *CCA1* and *TOC1* (Figure 5). TPC20 protein binds to the *CCA1* promoter in the *lwd1 lwd2* mutant and TT8 and TT2 can bind to target promoters in the *ttg1* mutant(17, 44), indicating that TLWD is not required for TF binding to DNA. However, in the absence of TLWD function these transcription factors cannot activate target gene expression. Similarly, we found a common defect of transcriptional repression of PRR-direct targets such as *CCA1*, *PIF4* and *PIF5* in higher order *prr* and *tlwd* mutants, indicating that TLWD is required for PRR-mediated repression. The essential function of TLWD might be due to simultaneous recruitment of additional transcriptional regulators and regulatory proteins. For example, it was reported that LWD1 forms complexes with YET ANOTHER KINASE (YAK1) and that *yak1* mutants have circadian phenotypes(46). More recently, LWD1 was implicated in the phase separation of CRY2-TCP22-PKK complexes. Understanding the composition and regulated assembly and disassembly of TLWD-clock complexes throughout the diel cycle might uncover novel targets to alter crop yield components such as growth and flowering time. More broadly, TLWD can be used as a model of the interplay between repression and activation required for dynamic transcriptional regulation.

### A new dynamic in circadian regulation

Rhythm generation in circadian biology has in recent times been considered analogous to a “clock”(47). Whilst clock analogies have their limitations(48), they have been useful in understanding the mechanistic basis of circadian timing. The circadian clocks are thought to comprise oscillators of transcriptional translational feedback loops (TTL) and separate or coupled feedback loops of metabolism and signalling(48, 49). The Arabidopsis TTL comprises the sequential expression of CCA1/LHY-PRRs-EC transcriptional repressors from dawn to end of night, and TCP and RVE8 transcriptional activators active at dawn and at midday, respectively(17, 50, 51). In the clock model the sequential expression and repression of the transcriptional activators are considered analogous to the wheels or cogs of a mechanical clock(48, 52). Essential to the sequential activities of the transcriptional repressors and activators are rhythmic changes in the abundance of the encoding transcripts and the proteins(13, 33, 50, 53). Here we find that TLWDs contribute to circadian timing by forming complexes with both transcriptional repressors and activators. TLWD proteins are present throughout the circadian cycle and do not appear to undergo large oscillatory changes in abundance, allowing for interactions with their binding partners as those change in abundance through the cycle (Figure 5). Thus, TLWD proteins represent a new category of circadian oscillator component, that are continuously present and essential for the function of the moving parts. In the clock analogy, the TLWD proteins act as arbors or spindles, being attached to the gears or wheels of the clock formed by the transcriptional repressors/activators. The arbor TLWD proteins provide the pivotal centre of the clock, attached to which the wheels of the clock can turn.

## Author contributions

A.A.R.W., E.H. and B.J.G. conceived the project and designed the experiments. EH conducted all the experiments. D.C.R. contributed to RNAseq analysis. E.H., B.J.G. and A.A.R.W. wrote the manuscript. All authors commented on the manuscript before submission.

## Acknowledgments

We thank M. Stancombe and M. Dorling for laboratory and plant growth support. We thank Prof. David Somers and Dr. Tina Schneider for providing guidance on co-immunoprecipitation experiments. We thank Prof. Antony Dodd for valuable critical comments on the manuscript. E.H. is supported by a Broodbank Fellowship from the Department of Plant Sciences at the University of Cambridge awarded to E.H., and a UK Space Agency International Bilateral Fund Pathfinder Phase 2 grant 10089325 awarded to A.A.R.W. Research performed by E.H. was also supported by a Leverhulme Trust Award RPG-2018-318 awarded to A.A.R.W and B.J.G. D.C.R. was supported by UKRI BBSRC grant BB/S002251/1 awarded to A.A.R.W.

## Data availability

All data that support the findings of this study are available at the Apollo repository (https://doi.org/10.17863/CAM.115480). RNAseq Raw data (FASTQ) files has been deposited under BioProject PRJNA1217148.

## Methods

### Plant materials and growth conditions

See Table S2 for details of the *A. thaliana* lines used in this study. Seedlings were surface sterilized by sequential immersion in 100% ethanol, 33% bleach with 0.01% (v/v) Triton X-100 and ultra-pure sterile water. Seeds were plated on Murashige and Skoog (MS) media (Duchefa) supplemented with 0.8% of bactoagar. After 3 days of stratification at 4 °C, they were placed in a controlled cabinet with constant 20 °C and different photoperiods of white light (130 μmol□m^−2^□s^−1^).

### Generation of transgenic lines

*Col-0 ttg1 lwd1 lwd2 pCCA1::LUC* were already described(27). Col-0 *ttg1 lwd1 lwd2* plants were crossed with either Col-O *pPRR7::LUC*(54) or Col-0 *pTOC1::LUC*(55). F1 heterozygous plants were backcrossed with *ttg1 lwd1 lwd2*, and *ttg1 lwd1 lwd2* with the relevant LUC reporter were selected in the next generation. Mutants were confirmed by genotyping using previously described primers(27). In the next generation, homozygous LUC reporter was confirmed by lack of segregation for resistance to phosphinothricin (PPT) for *pPRR7::LUC*, or hygromycin (HYG) for *pTOC1::LUC* when seedlings were grown on agar supplements with the selection media.

For LWD1-Venus and Venus-TTG1 complementation lines, LWD1 promoter (1068 bp) or TTG1 promoter (1328bp), LWD1 or TTG1 coding sequence, mVenus, downstream sequences (dss) of LWD1 (713 bp) or TTG1 (1000bp) were cloned in the pGREEN-BASTA binary vector by Gibson assembly to obtain pLWD1::LWD1-Venus::LWD1dss and pTTG1::Venus-TTG1::TTG1dss. For LWD1-LUC+ and TTG1-LUC+ complementation lines, the plasmids pCsA-BASTA-pLWD1::LWD1-LUC::LWD1dss or pCsA-BASTA-pTTG1::TTG1-LUC+::TTG1-dss were obtained by loop assembly(56). A flexible linker was inserted between the coding sequence of LWD1 and TTG1 and Venus or LUC+. The corresponding plasmids were transformed by floral dipping into *lwd1 lwd2 pCCA1::LUC* or *ttg1-21 pCCA1::LUC*. T0 seeds were selected on MS PPT plates. In T1, at least two independent lines were selected on their ability to complement mutant phenotypes: circadian periodicity of *pCCA1::LUC* and flowering time for *lwd1 lwd2*, and seed coat pigmentation and trichomes for *ttg1-21*. Then T2 homozygous lines were obtained from selected lines and the presence of the fusion protein was confirmed by western blots in either T2 or T3 seedlings.

### Measuring diel and circadian rhythms of LUC reporter lines

Luminescence assays were performed as described in (27). Clusters of 5 plants with the indicated LUC reporters were grown for 10 days under entrainment photoperiods before luciferin solution was added to the media and plants were placed in the imaging system (either a Photek ICCD25 or a Berthold camera system). Every hour, bioluminescence signal was integrated for 800 seconds. Similar light intensity entrainment conditions were maintained for 2-3 days, followed by 3-5 days of continuous light. Where indicated, traces were normalized to the average bioluminescence through the experiment.

### RNAseq

Seeds were grown for 10 days in ½MS plates supplemented with 0.8% agar under diel cycles of 12 light / 12 dark. Seedlings were harvested at time ZT6 and time ZT18 and immediately frozen in liquid nitrogen. Each technical replicate consisted of three clusters of 10 seedlings growing on different agar plates in the same controlled cabinet. Plant material was ground using a Tissue Lyser (QIAGEN). Total RNA was extracted with the RNeasy Plant Mini Kit (QIAGEN). After extraction, the quality of the RNA was determined with a Bioanalyzer (Agilent Technologies). Three biological replicates were obtained for each time point, by pooling the RNA of three technical replicates. Library preparation and pair-end sequencing was performed with the Illumina RNA-seq platform with 15-20 million 150bp-reads by Genewiz. Sequencing quality was assessed by generating FASTQ(57). Trimmomatic(58) was used to remove adapters and non-Arabidopsis sequences. We used RNA STAR(59) to map sequences to the Arabidopsis genome using the TAIR10 assembly (Arabidopsis_thaliana.TAIR10.49.gff3). Mapped reads were quantified with featureCounts(60). Differential gene expression between Col-0 and *ttg1*, *lwd1 lwd2* and *ttg1 lwd1 lwd2* mutants was calculated with DESeq2(61). Genes with a fold change Log2 ≥ 1.5 with respect to wildtype were considered DEGs. We used the phase enrichment tool in the CAST-R database(30) to analyse circadian phase of selected DEGs using Romanowski et al_LL_LDHH(4) as the reference dataset.

### qPCR

Seedlings were grown for 10 days under short day photoperiods (8h light /16h dark). For each time point, three biological replicates (each formed by three pooled clusters of approximately 20 seedlings) were collected, frozen in liquid nitrogen and ground with a Tissue Lyser (QIAGEN). Total RNA was extracted using the RNeasy Plant Mini Kit (QIAGEN) and RNase-Free DNase on-column treatment (QIAGEN). cDNA was synthesized from 500□ng RNA with the RevertAid First Strand cDNA Synthesis Kit (Thermo Scientific) using oligo (dT) primers. The gene-specific products were amplified using the Quantinova SYBR Green PCR Kit (QIAGEN) on a Biorad CFX384 Real-Time System. Primers used are detailed in Supplementary Table S3. Relative transcript levels were normalized to both *PP2A* and *UBQ10* expression.

### Yeast 2-hybrid experiments

All coding sequences were amplified from Col-O cDNA and cloned into pDONR201 by Gateway BP reaction (Invitrogen) and then cloned into various yeast expression vectors by LR reaction (Invitrogen). LWD1, LWD2 and TTG1 were cloned into both pAS2 (Clontech) and pDEST 32 (Invitrogen) to generate N-terminal GAL4-BD fusion proteins. PRR9, PRR7, PRR5, TOC1, CCA1, LHY, RVE1 and RVE8 coding sequences were cloned into pC-ACT2 (Clontech) and TCP20, TCP21 and TCP22 were cloned into pDEST22 to generate N-terminal GAL4-BD fusion proteins. Plasmids were verified by Sanger sequencing. AH109 were transform with the GAL4-BD vectors and Y187 and Y187 cells were transformed with GAL4-BD vectors, by Li-Ac mediated transformation following Clontech Yeast Protocols Handbook (Clontech). Positive colonies were re-streaked in the appropriate synthetic dropout (SD) media lacking leucine (-L) or tryptophane (-W). Appropriate pAS+pACT2 or pDEST32+pDEST22 diploid strains were obtained by mating and selecting in SD-LW media. Protein interactions were evaluated by the ability of diploid strains to grow on SD media without Histidine and Adenine. 3-amino-1,2,4-triazole (3AT) was added to increase the stringency of Histidine selection. Serial dilutions of diploid strains were plated in SD - LW, SD-LHW supplemented with (3AT) and SD-LHWA and grown for 3-4 days before imaging. Y2H results were confirmed in at least 2 independent experiments.

### Arabidopsis whole protein extracts and western blot

Approximately 40 15-day old seedlings were collected and frozen in nitrogen at specific times of the diel cycle. Plant materials were finely ground in a Tissue lyser and resuspended in fresh extraction buffer (50 mM Tis-HCl pH 7.5, 150 mM Na CL, 0.5% NP-40, 1mM EDTA, 2mM Na_3_VO_4_, 10 mM NaF, 2mM β-glycerol phosphate, 1mM PMSF, 50 μM MG132, 50 μM MG115, 50 μM ALLN, 3mM DTT and Protein inhibitor tablet). Protein extracts were cleared by centrifugation at max speed at 4°C. Protein concentration was measured with Pierce™ 660nm Protein Assay Reagent (Thermo Scientific) following the manufacturer’s instructions. 6X SDS buffer was added to protein extracts. 10 μg of each sample was loaded into precast gels (SurePAGE Bis-Tris, GenScript). Proteins were transferred onto supported nitrocellulose membrane (Biorad) by wet transfer. Membrane was blocked with Western blocking reagent (Roche) and incubated with α-GFP (Roche, 1181446000) and α-mouse-HRP (Proteintech SA00001-1). Membranes were covered with ECL SuperBright (Agrisera) for 2 minutes and briefly rinsed in TBS before imaging with a G:BOX Chemi XRQ imaging system or an Amersham™ ImageQuant™ 800 biomolecular imager.

### Co-Immunoprecipitation

The coding sequences of LWD1 and TTG1 were cloned into pCK1_35S::mTurq-LWD1 and pCK1_p35S::mTurq-TTG1 by Loop assembly cloning(62). The coding sequences of PRR9, PRR7, PRR5 and TOC1 were cloned into pJCV52(63) to obtain pJCV52_p35S::PRR-HA by Gateway cloning. Plasmids were introduced into Agrobacterium strain AGL1. Appropriate Agrobacterium cultures were diluted and mixed with p19 plasmid to OD_600_=0.5 in infiltration solution (1/4MS pH 6, 1% sucrose, 100μM acetosyringone and 50 μl/l Silwet L-77) and left for 3-4 hours in the dark before infiltrating leaves of 3-4 week old *Nicotiana benthamiana* plants with a 1mL syringe. Plants were kept in the light until leaves were dried, then covered with a black bag for 24 hours. Infiltrated leaves were collected 48-72 hours after infiltration and immediately frozen in liquid nitrogen. Plant material was finely ground in a Tissue lyser and resuspended in fresh extraction buffer(64) (50 mM Tis-HCl pH 7.5, 150 mM Na CL, 0.5% NP-40, 1mM EDTA, 2mM Na_3_VO_4_, 10 mM NaF, 2mM β-glycerol phosphate, 1mM PMSF, 50 μM MG132, 50 μM MG115, 50 μM ALLN, 3mM DTT and Protein inhibitor tablet). Protein extracts were cleared by centrifugation at max speed at 4°C then incubated with µMACS^TM^ HA beads (Miltenyi Biotec) for 2 hours with rotation at 4°C. Columns (Miltenyi Biotec) were used to wash the beads with extraction buffer following the manufacturer’s protocol. To elute protein complexes, HA beads were incubated for 15 minutes with HA peptide solution at 30°C. Then, 6X sample buffer was added to the eluate and incubated for 1 min at 90°C before loading into precast gels (SurePAGE Bis-Tris, GenScript). Electrophoresis was performed at 4°C and low voltage (60-90V). Proteins were transferred onto supported nitrocellulose membrane (Biorad) by wet transfer at 4°C. Membrane was blocked with Western blocking reagent (Roche) and incubated with primary antibodies α-HA (Abcam, ab9110) and α-GFP (Roche, 1181446000) and secondary antibodies α-rabbit-HRP (Proteintech SA00001-2) and α-mouse-HRP (Proteintech SA00001-1), respectively. Membranes were covered with ECL SuperBright (Agrisera) for 2 minutes and briefly rinsed in TBS before imaging with a G:BOX Chemi XRQ imaging system.

### Hypocotyl elongation

Seeds were plated on MS media (4.4 g/L with 0.5 g/L MES and 1.5% phytoagar pH 5.7). After stratification plants were placed in a growth cabinet with 100 μmol□m^−2^□s^−1^ of white light under short day (8h_L/ 16D) or long day (16h_L/ 8h_D) photoperiods. Hypocotyl length was measured after 7 days using Image J software as previously described(65).

## Supplementary Tables

**Table S1. RNAseq differentially expressed genes Log2^FC^, normalized counts and circadian phase of expression**

**Table S2. *Arabidopsis thaliana* lines used in this study**

**Table S3. qPCR primers**

## Supplementary Figure Legends

**Figure S1.**
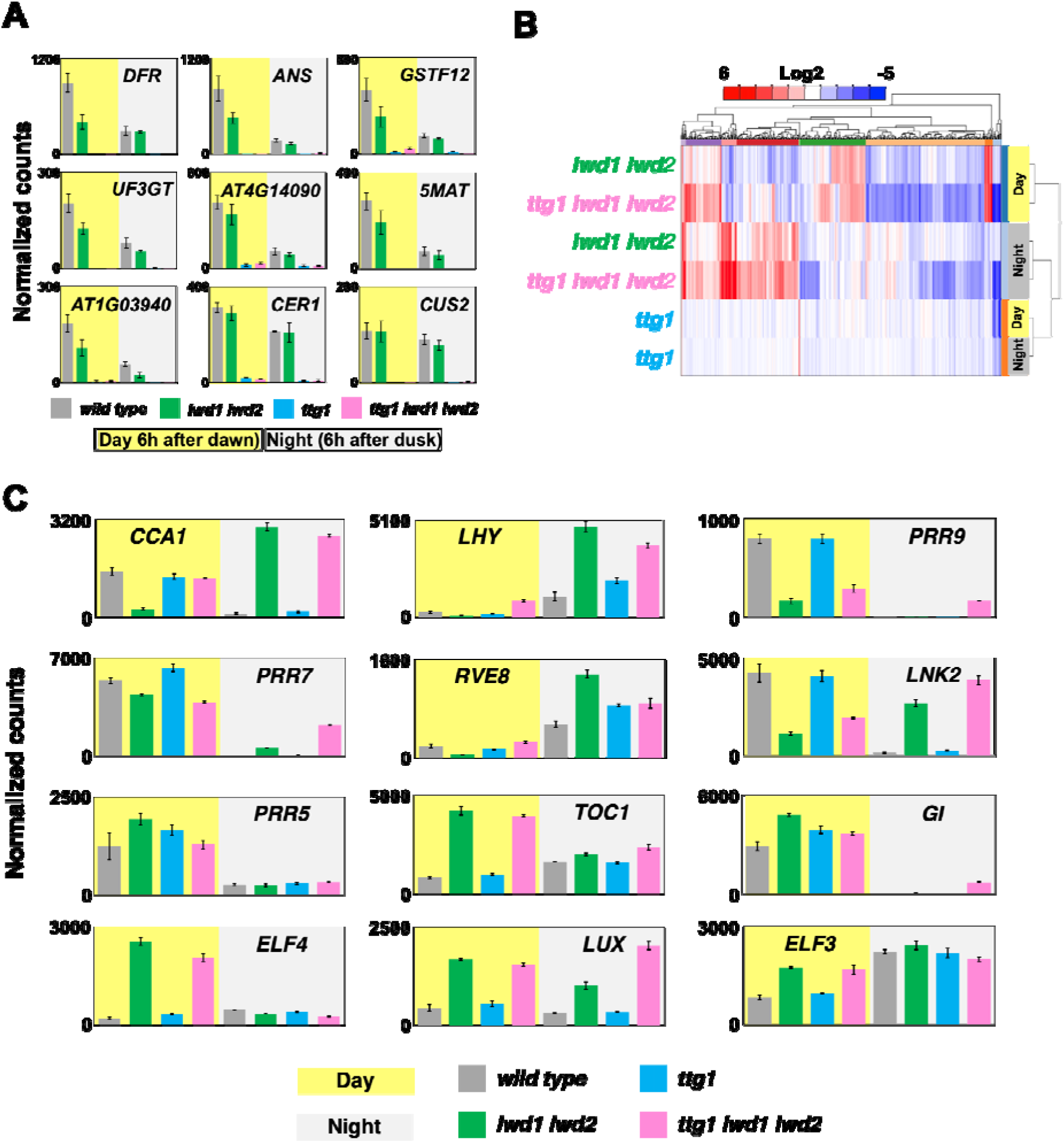
Transcript abundance of MBW complex targets and circadian oscillator components in *tlwd* mutants. (A) Normalized expression of MBW complex direct targets^29^ in wild type, *lwd1 lwd2*, *ttg1* and *ttg1 lwd1 lwd2* mutants. (B) Hierarchical clustering of DEGs in *ttg1*, *lwd1 lwd2* and *ttg1 lwd1 lwd2* at Day (ZT6) and Night (ZT18) when compared to wild type expression. (C) Normalized expression of selected circadian oscillator components in wild type, *lwd1 lwd2*, *ttg1* and *ttg1 lwd1 lwd2* mutants. Normalized counts in A and C were obtained with Deseq2. Error bars depict SEM, n=3 See Table S1 for values.

**Figure S2.**
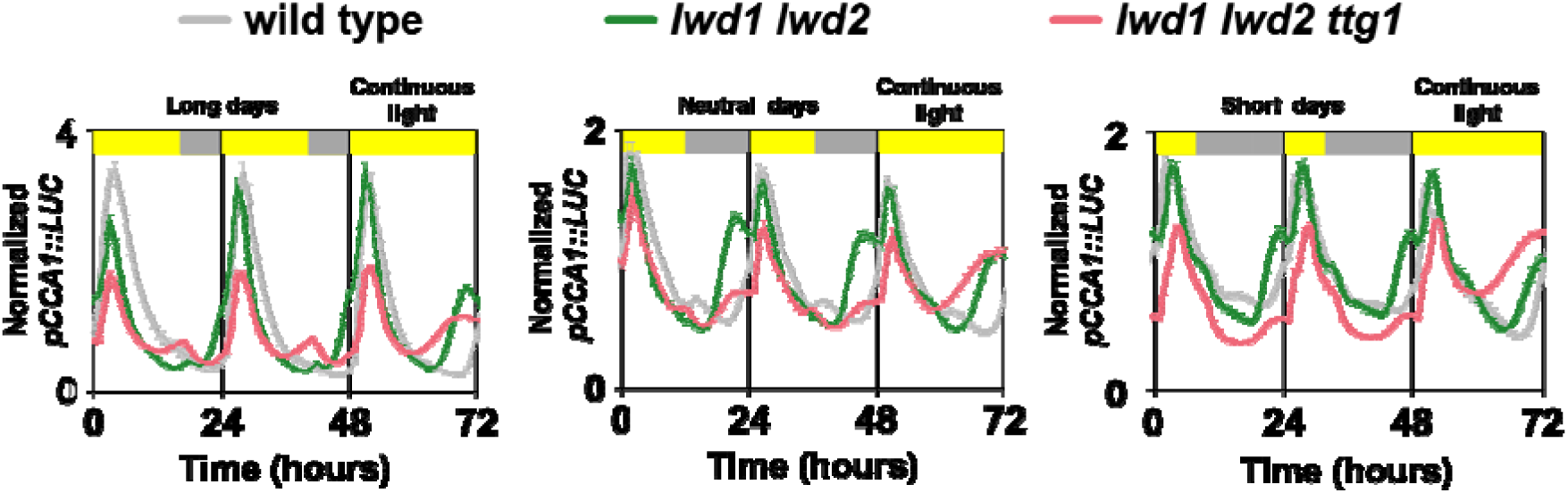
Rhythms of *pCCA1:LUC* under different photoperiods in wild type, *lwd1 lwd2* and *ttg1 lwd1 lwd2* mutants. Note that under long days (left panel) the *ttg1 lwd1 lwd2* mutant has a second peak of *CCA1* in the second half of the diel cycle which is not apparent under neutral days or short days (middle and right panels). Error bars depict SEM, n= (LD) wild type (8), *lwd1 lwd2* (16) and *ttg1 lwd1 lwd2* (12); (ND) wild type (12), *lwd1 lwd2* (12) and *ttg1 lwd1 lwd2* (12); (SD) wild type (24), *lwd1 lwd2* (22) and *ttg1 lwd1 lwd2* (28).

**Figure S3.**
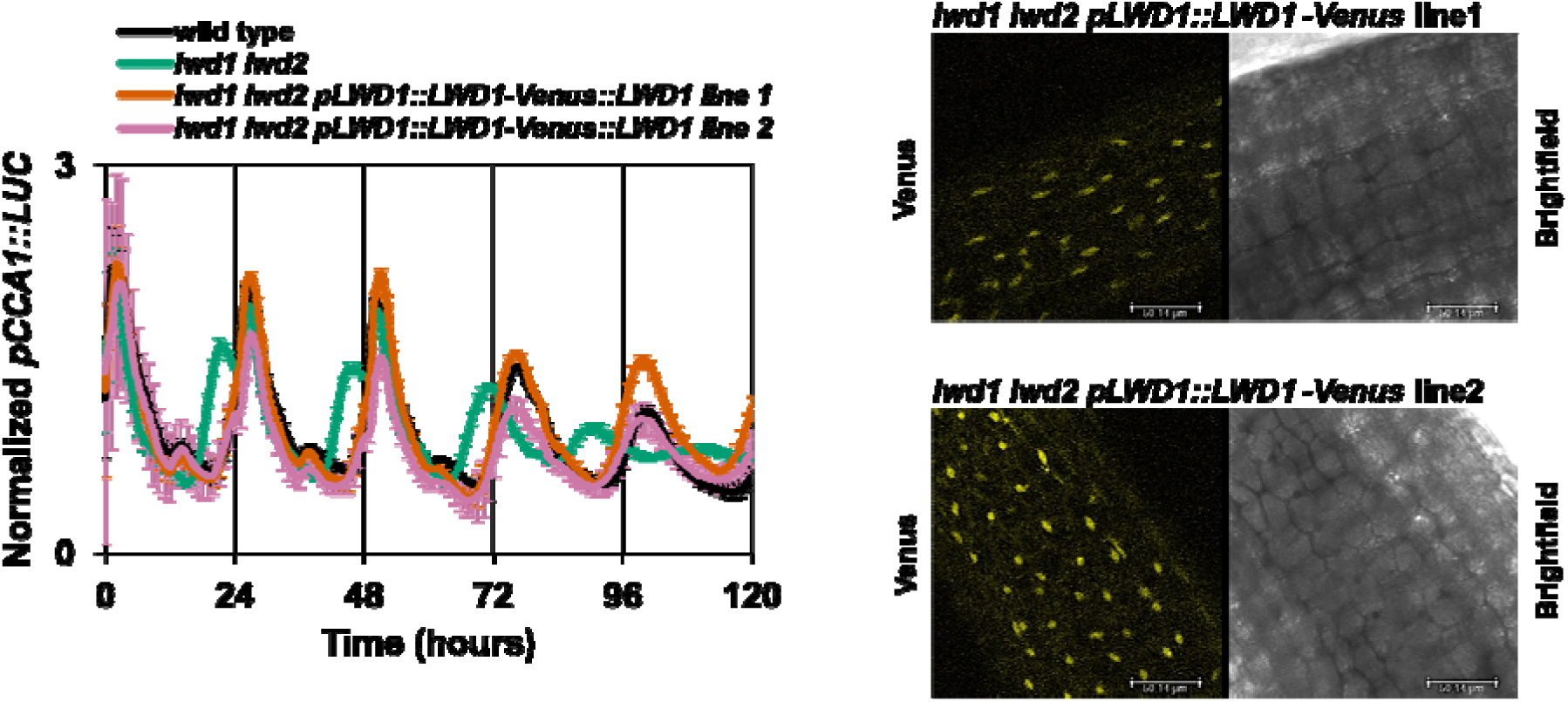
*LWD1-Venus* complementation lines. (A) Complementation of *lwd1 lwd2* mutant short periodicity phenotype in two independent *pLWD1::LWD1-Venus::LWD1dss lines*. Y-axis depict average normalized bioluminescence, error bars depict SEM, n= wild type (12), *lwd1 lwd2* (12), *lwd1 lwd2 pLWD1::LWD1-Venus::LWD1* line 1(8) and line 2 (8). Confocal microscopy images show nuclear localization of LWD1-Venus in both complementation lines.

**Figure S4.**
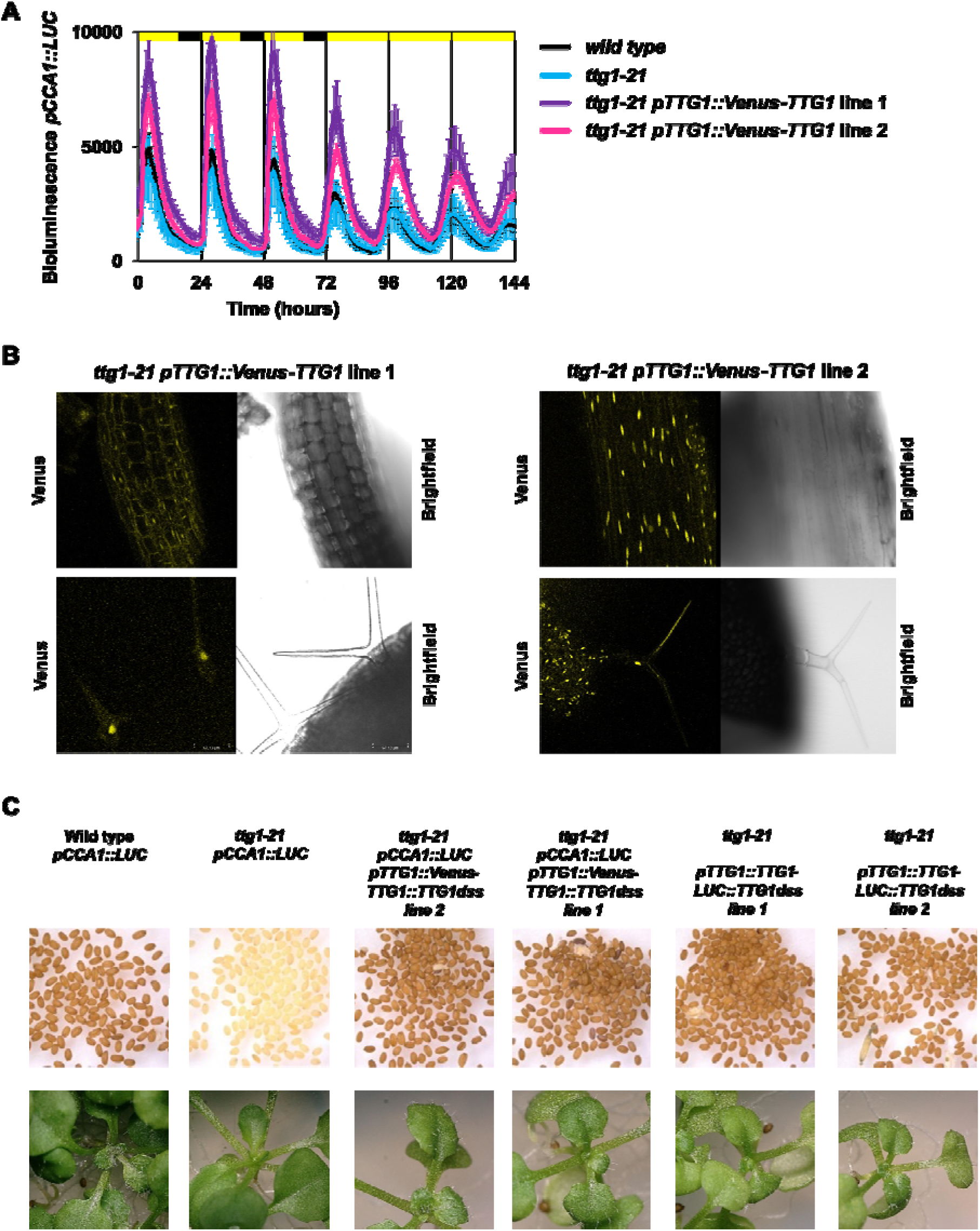
TTG1 complementation lines. (A) Oscillations of *pCCA1::LUC.* Complementation of *ttg1* mutant with *pTTG1::Venus-TTG1::TTG1dss* does not affect circadian rhythms. Note that *ttg1* does not have a circadian phenotype. Y-axis depicts average bioluminescence of 8 seedling clusters per genotype, error bars depict SEM. (B) Confocal microscopy images show nuclear localization of Venus-TTG1-Venus in complementation lines both in stem and trichomes. (C) Complementation of *ttg1-21* mutant lack of seed coat pigmentation and trichomes in *Venus-TTG1* and *TTG1-LUC* complementation lines.

**Figure S5.**
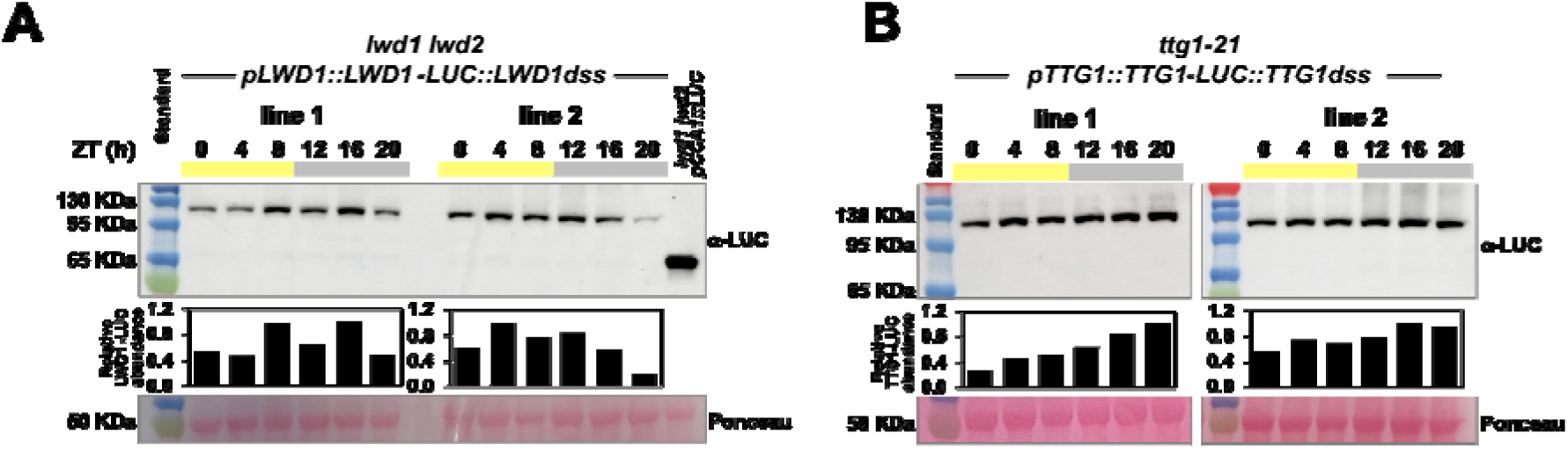
TLWD proteins are present throughout the diel cycle. Representative western blots from protein extracts of (A) *lwd1 lwd2 pLWD1::LWD1-LUC* and (B) *ttg1 pTTG1::Venus-TTG1*. Seedlings from two independent lines (Line 1 and Line 2) were grown for 10 days under diel cycles (12hL/12hD) and harvested at the indicated time points. Expected molecular weight for both LWD1-LUC and TTG1-LUC is 100 KDa. Bar graphs show protein abundance relative to maximum. Ponceau staining shown loading control, 10 µg of protein extract was loaded for each timepoint.

**Figure S6.**
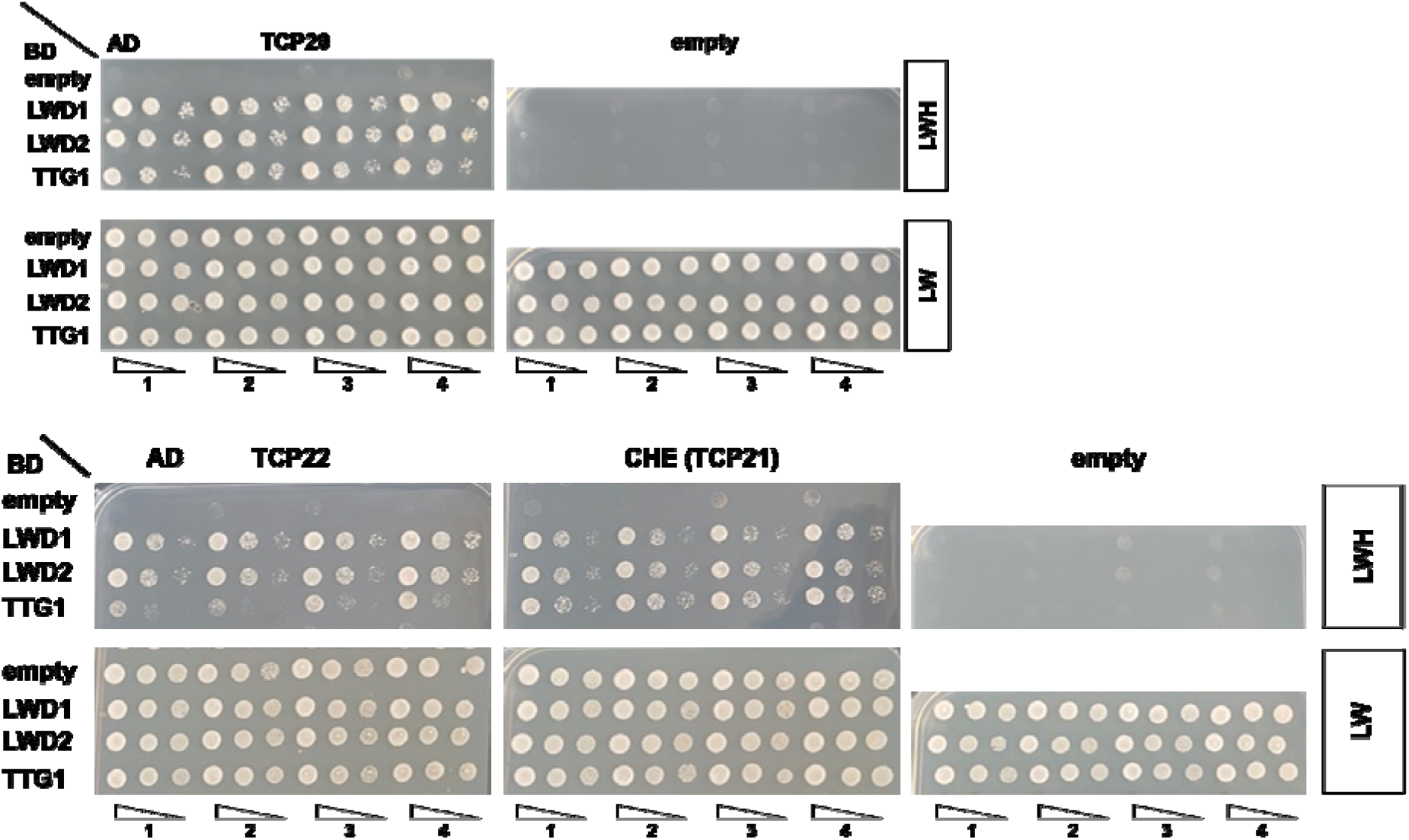
Interaction of TLWD proteins with TCPs. All three TLWD proteins interact with TCP20, TCP22 and CHE (TCP21). 3 serial dilutions (1:10) of four independent colonies. Growth in LHW media indicates interaction between protein pairs.

**Figure S7.**
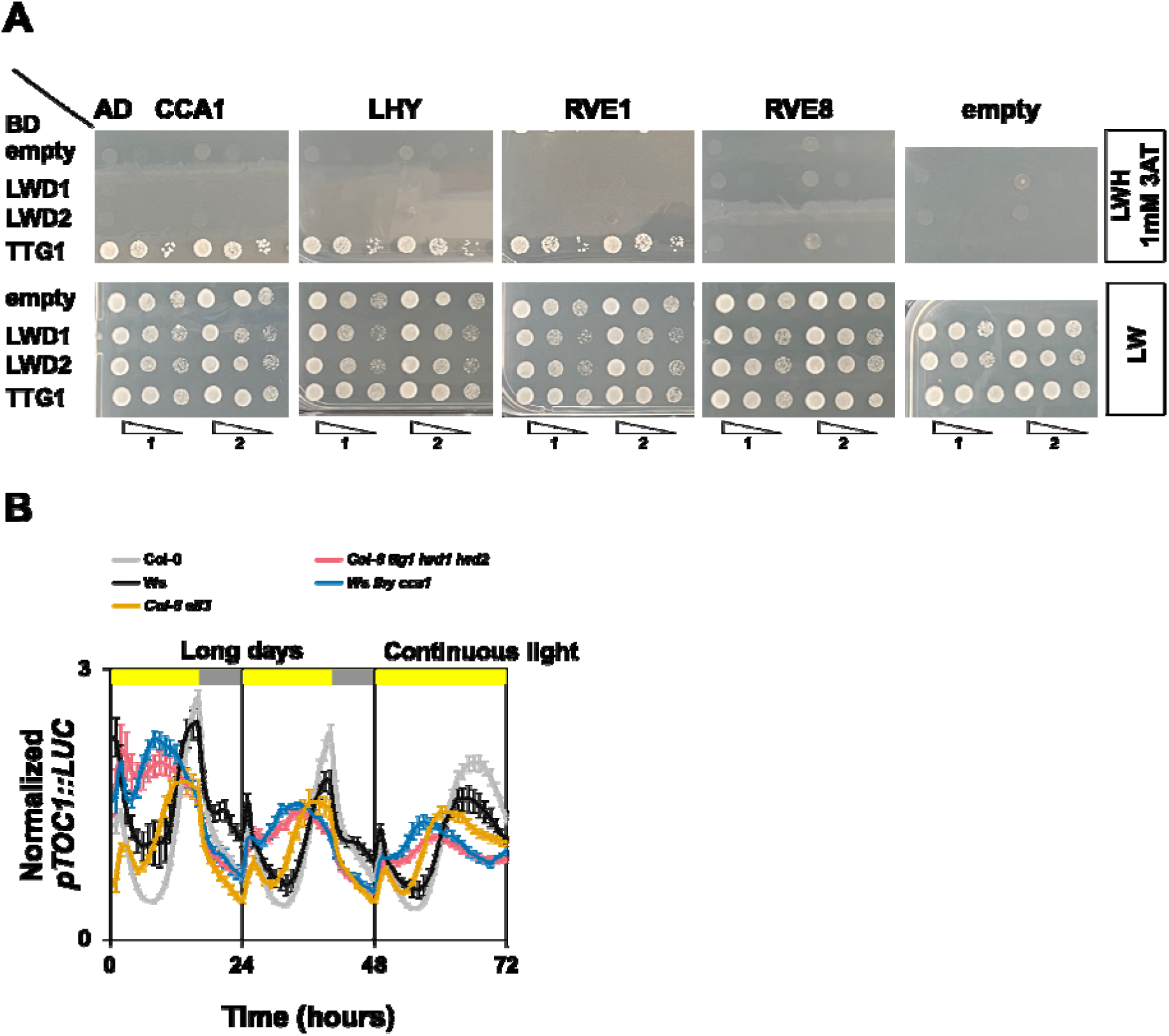
TLWD proteins interact with Myb-like transcription factors and control *TOC1* expression. (A) TTG1 interacts with CCA1, LHY and RVE1 but not with RVE8. 3 serial dilutions (1:10) of two independent colonies. Growth in LHW media with 1mM 3AT indicates interaction between protein pair. (B) Oscillations of *pTOC1::LUC* in wild type, *ttg1 lwd1 lwd2, cca1 lhy and elf3* under diel cycles and continuous light. Note that *ttg1 lwd1 lwd2* and *elf3* are in Col-0 background whereas *lhy cca1* is in the Ws background. Col-0 and Ws wild types have similar rhythms. Y-axis depicts average normalized bioluminescence of 8 clusters of seedlings per genotype.

**Figure S8.**
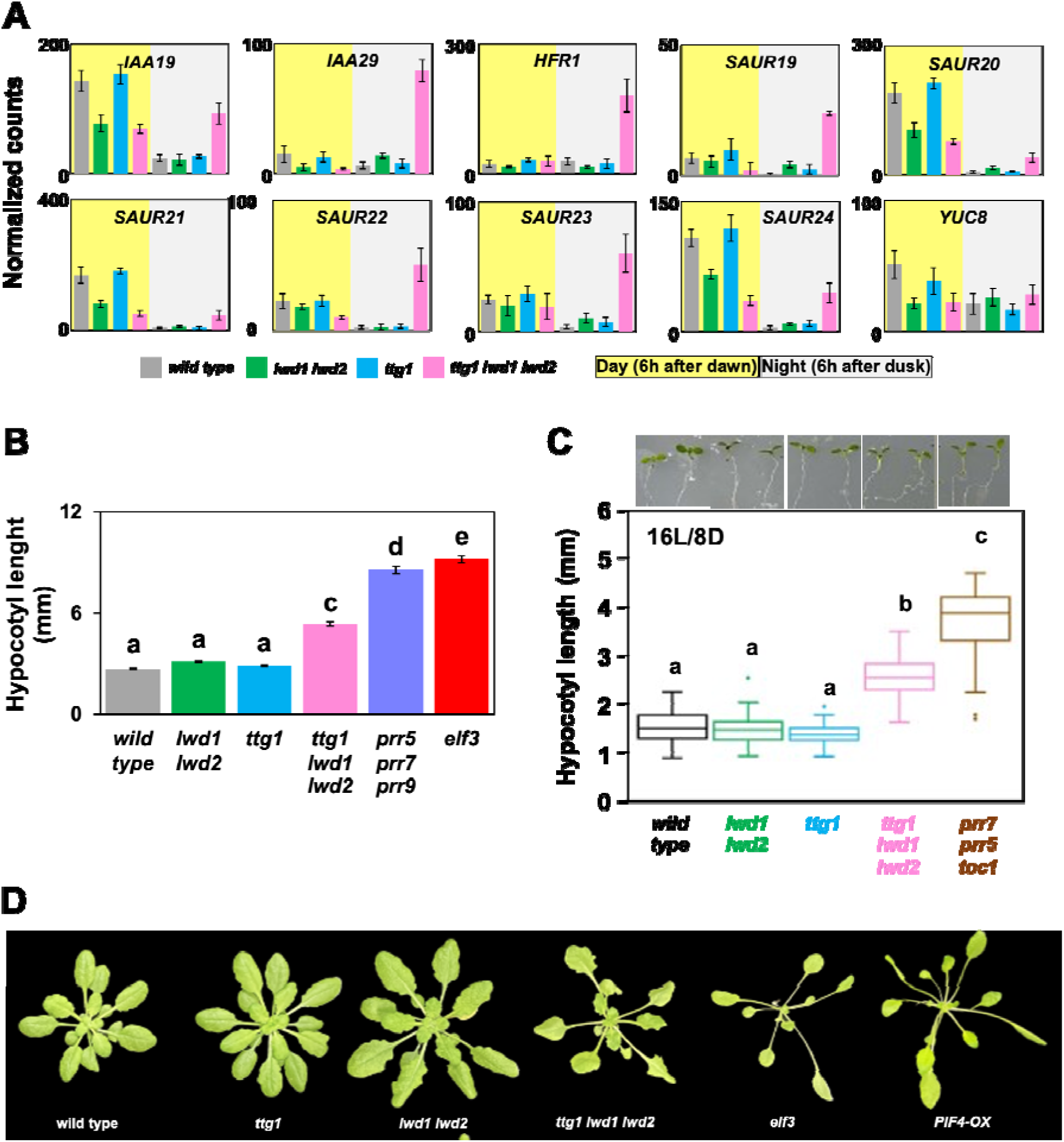
*ttg1 lwd1 lwd2* mutants recapitulate some *PIF4-OX* phenotypes. (A) Normalized expression of PIF4/5 targets in wild type and *ttg1 lwd1 lwd2* mutants. Error bars depict SEM, n=3. Normalized counts were obtained with Deseq2. See Table S1 for values. B) Hypocotyl growth of plants grown for 7 days under SD. Error bars depict SEM, n= wild type (88), *ttg1* (91), *lwd1 lwd2* (73), *ttg1 lwd1 lwd2* (116), *prr5 prr7 prr9* (82) and *elf3* (103). Letters depict genotypes with statistically different average length with the Tukey Kramer test. C) Hypocotyl growth and representative images of seedlings grown for 7 days under LD. Error bars depict SEM, n= wild type (64), *lwd1 lwd2* (46), *ttg1* (71), *ttg1 lwd1 lwd2* (77) and *prr5 prr7 toc1* (48). (D) Representative rosettes of wild type, *ttg1*, *lwd1 lwd2*, *ttg1 lwd1 lwd2*, *elf3* and *PIF4-OX*(*66*) mutants grown under SD.

