## Supplementary material for "Arbor function of TRANSPARENT TESTA GLABRA 1 and LIGHT REGULATED WD scaffold proteins in the Arabidopsis circadian oscillators includes transcriptional repression through PSEUDO RESPONSE REGULATORS": Figure S1

### Slide 1
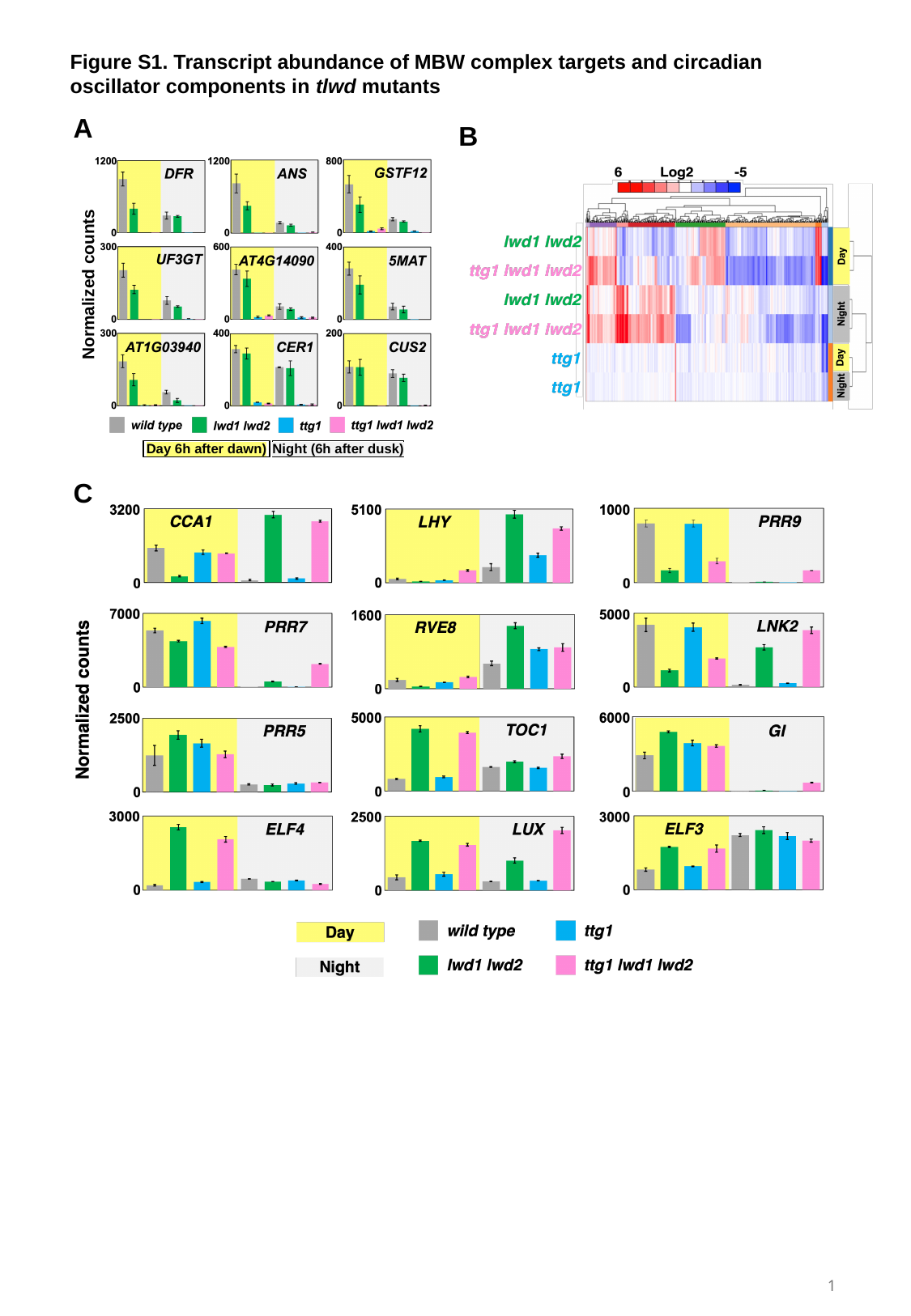

Figure S1. Transcript abundance of MBW complex targets and circadian oscillator components in tlwd mutants
A
B
Normalized counts
Day 6h after dawn)
Night (6h after dusk)
C
1
