## Supplementary material for "Arbor function of TRANSPARENT TESTA GLABRA 1 and LIGHT REGULATED WD scaffold proteins in the Arabidopsis circadian oscillators includes transcriptional repression through PSEUDO RESPONSE REGULATORS": Figure S2

### Slide 1
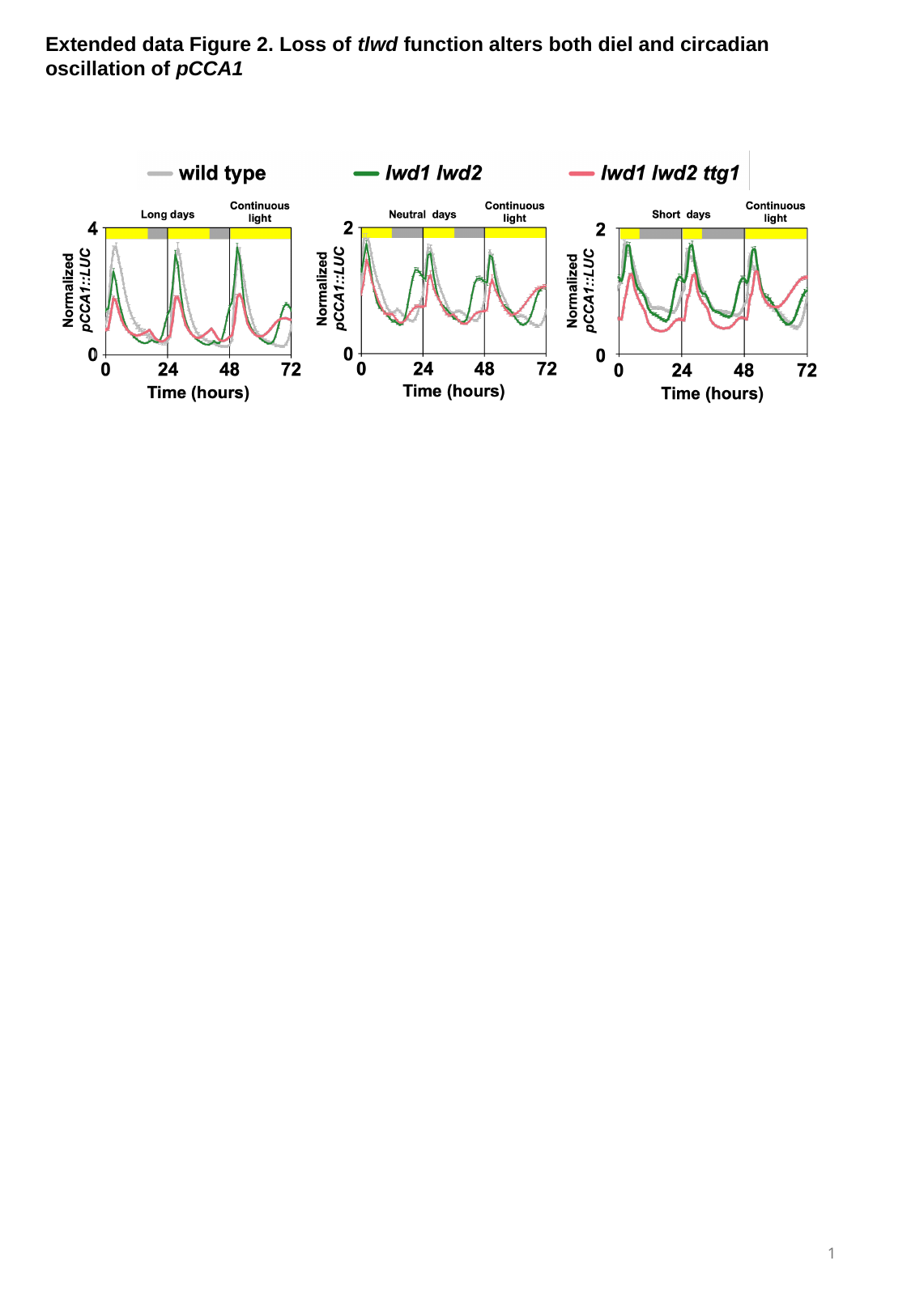

Extended data Figure 2. Loss of tlwd function alters both diel and circadian oscillation of pCCA1
1
