## Supplementary material for "Arbor function of TRANSPARENT TESTA GLABRA 1 and LIGHT REGULATED WD scaffold proteins in the Arabidopsis circadian oscillators includes transcriptional repression through PSEUDO RESPONSE REGULATORS": Figure S3

### Slide 1
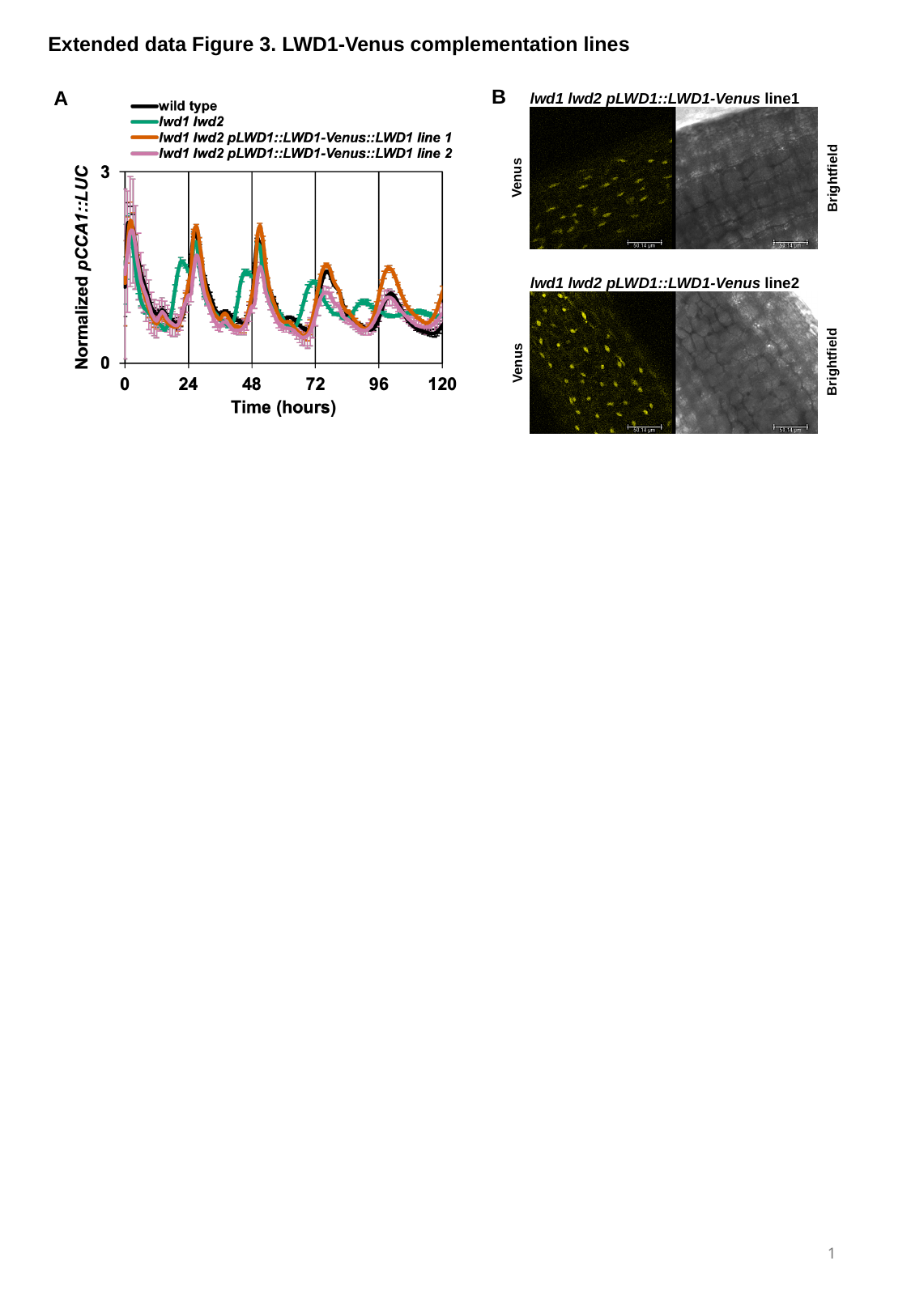

Extended data Figure 3. LWD1-Venus complementation lines
B
A
lwd1 lwd2 pLWD1::LWD1-Venus line1
Venus
Brightfield
lwd1 lwd2 pLWD1::LWD1-Venus line2
Brightfield
Venus
1
