## Supplementary material for "Arbor function of TRANSPARENT TESTA GLABRA 1 and LIGHT REGULATED WD scaffold proteins in the Arabidopsis circadian oscillators includes transcriptional repression through PSEUDO RESPONSE REGULATORS": Figure S4

### Slide 1
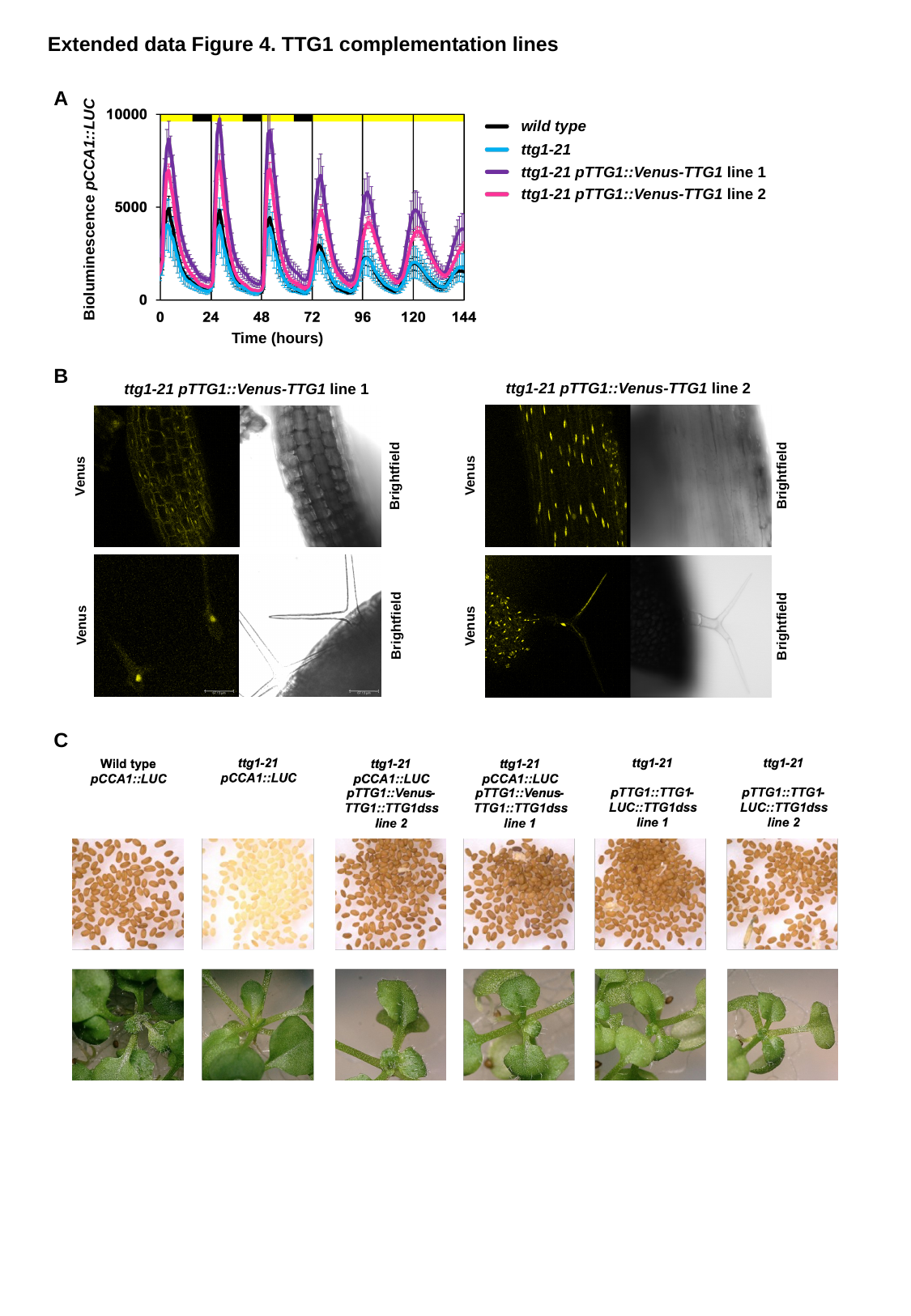

Extended data Figure 4. TTG1 complementation lines
A
wild type
ttg1-21
ttg1-21 pTTG1::Venus-TTG1 line 1
ttg1-21 pTTG1::Venus-TTG1 line 2
Bioluminescence pCCA1::LUC
Time (hours)
B
ttg1-21 pTTG1::Venus-TTG1 line 2
ttg1-21 pTTG1::Venus-TTG1 line 1
Venus
Brightfield
Venus
Brightfield
Venus
Brightfield
Venus
Brightfield
C
