## Supplementary material for "Arbor function of TRANSPARENT TESTA GLABRA 1 and LIGHT REGULATED WD scaffold proteins in the Arabidopsis circadian oscillators includes transcriptional repression through PSEUDO RESPONSE REGULATORS": Figure S5

### Slide 1
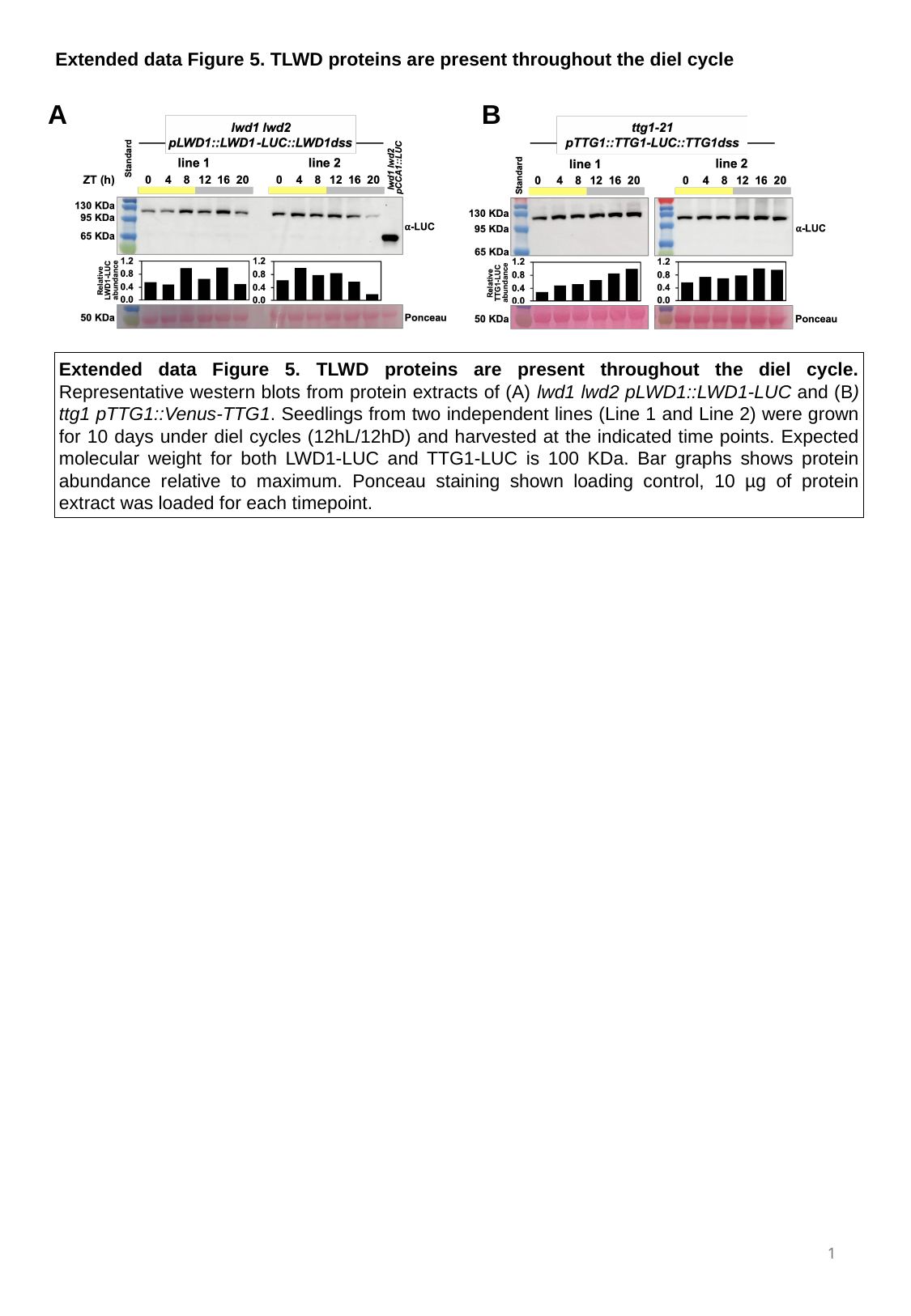

Extended data Figure 5. TLWD proteins are present throughout the diel cycle
A
B
Extended data Figure 5. TLWD proteins are present throughout the diel cycle. Representative western blots from protein extracts of (A) lwd1 lwd2 pLWD1::LWD1-LUC and (B) ttg1 pTTG1::Venus-TTG1. Seedlings from two independent lines (Line 1 and Line 2) were grown for 10 days under diel cycles (12hL/12hD) and harvested at the indicated time points. Expected molecular weight for both LWD1-LUC and TTG1-LUC is 100 KDa. Bar graphs shows protein abundance relative to maximum. Ponceau staining shown loading control, 10 µg of protein extract was loaded for each timepoint.
1
