## Supplementary figures and images for "Arbor function of TRANSPARENT TESTA GLABRA 1 and LIGHT REGULATED WD scaffold proteins in the Arabidopsis circadian oscillators includes transcriptional repression through PSEUDO RESPONSE REGULATORS"

### Figure S6

#### Slide 1
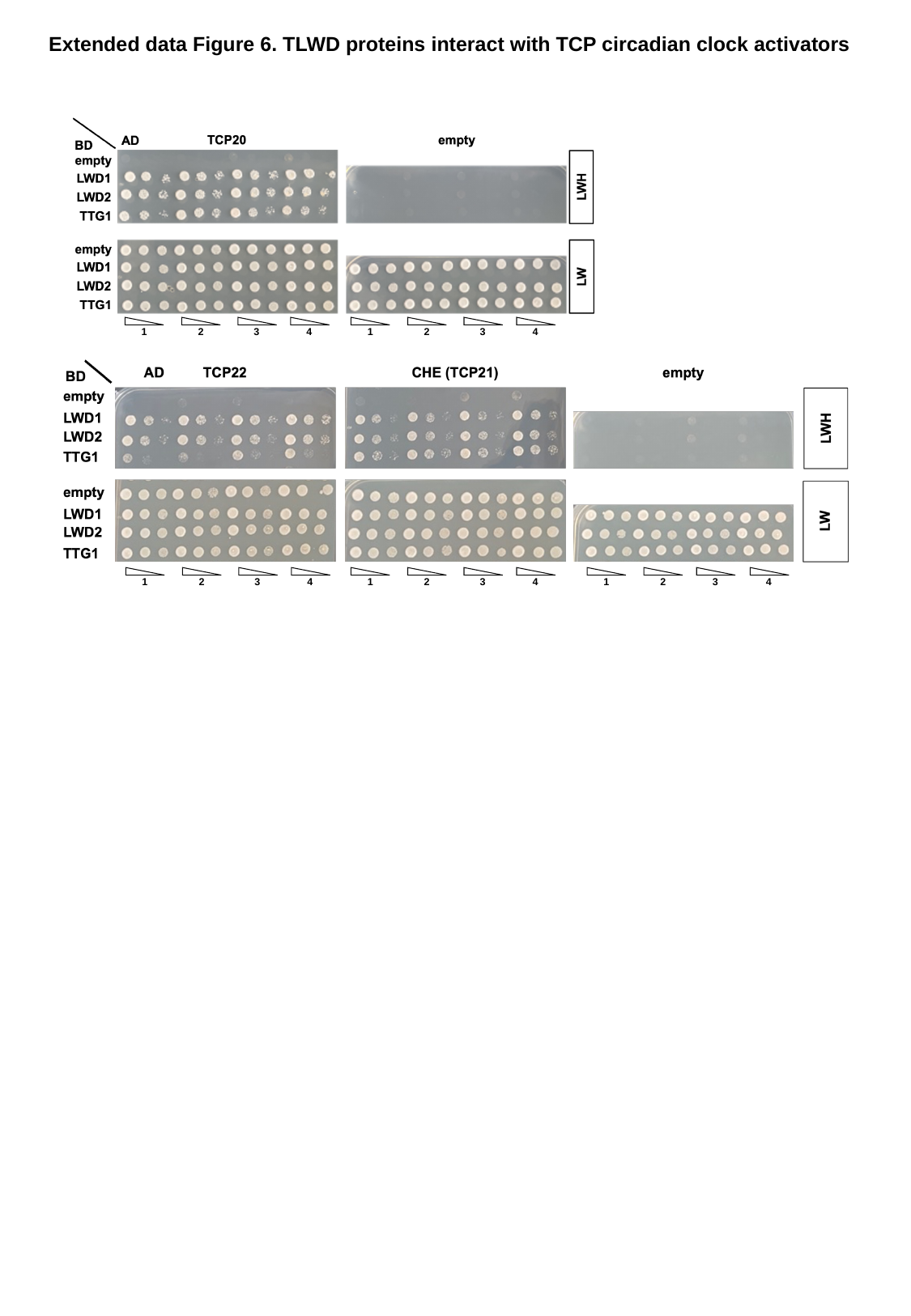

Extended data Figure 6. TLWD proteins interact with TCP circadian clock activators
1
2
3
4
1
2
3
4
1
2
3
4
1
2
3
4
1
2
3
4
