## Supplementary material for "Arbor function of TRANSPARENT TESTA GLABRA 1 and LIGHT REGULATED WD scaffold proteins in the Arabidopsis circadian oscillators includes transcriptional repression through PSEUDO RESPONSE REGULATORS": Figure S7

### Slide 1
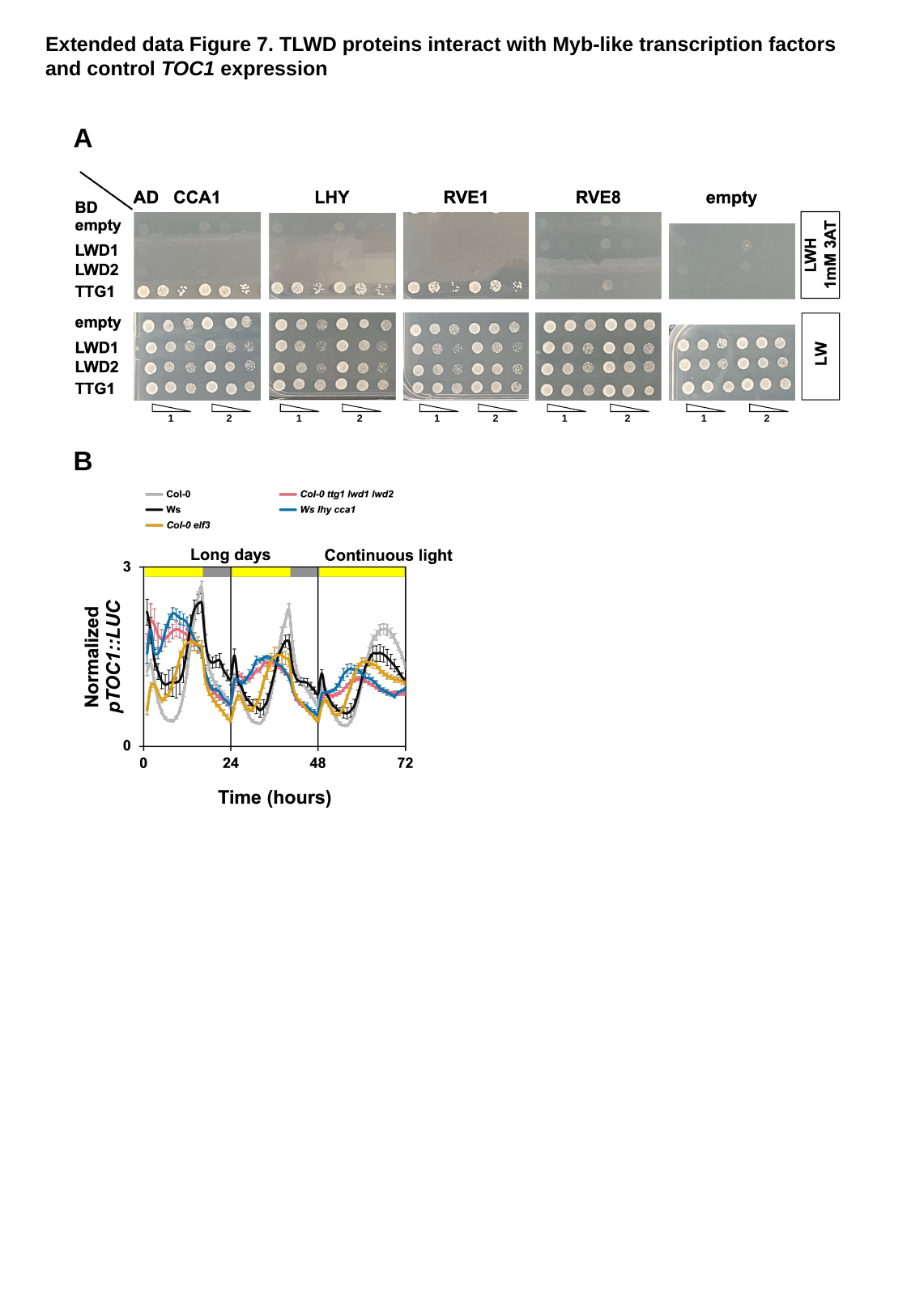

Extended data Figure 7. TLWD proteins interact with Myb-like transcription factors and control TOC1 expression
A
1
2
1
2
1
2
1
2
1
2
B
