## Supplementary material for "Arbor function of TRANSPARENT TESTA GLABRA 1 and LIGHT REGULATED WD scaffold proteins in the Arabidopsis circadian oscillators includes transcriptional repression through PSEUDO RESPONSE REGULATORS": Figure S8

### Slide 1
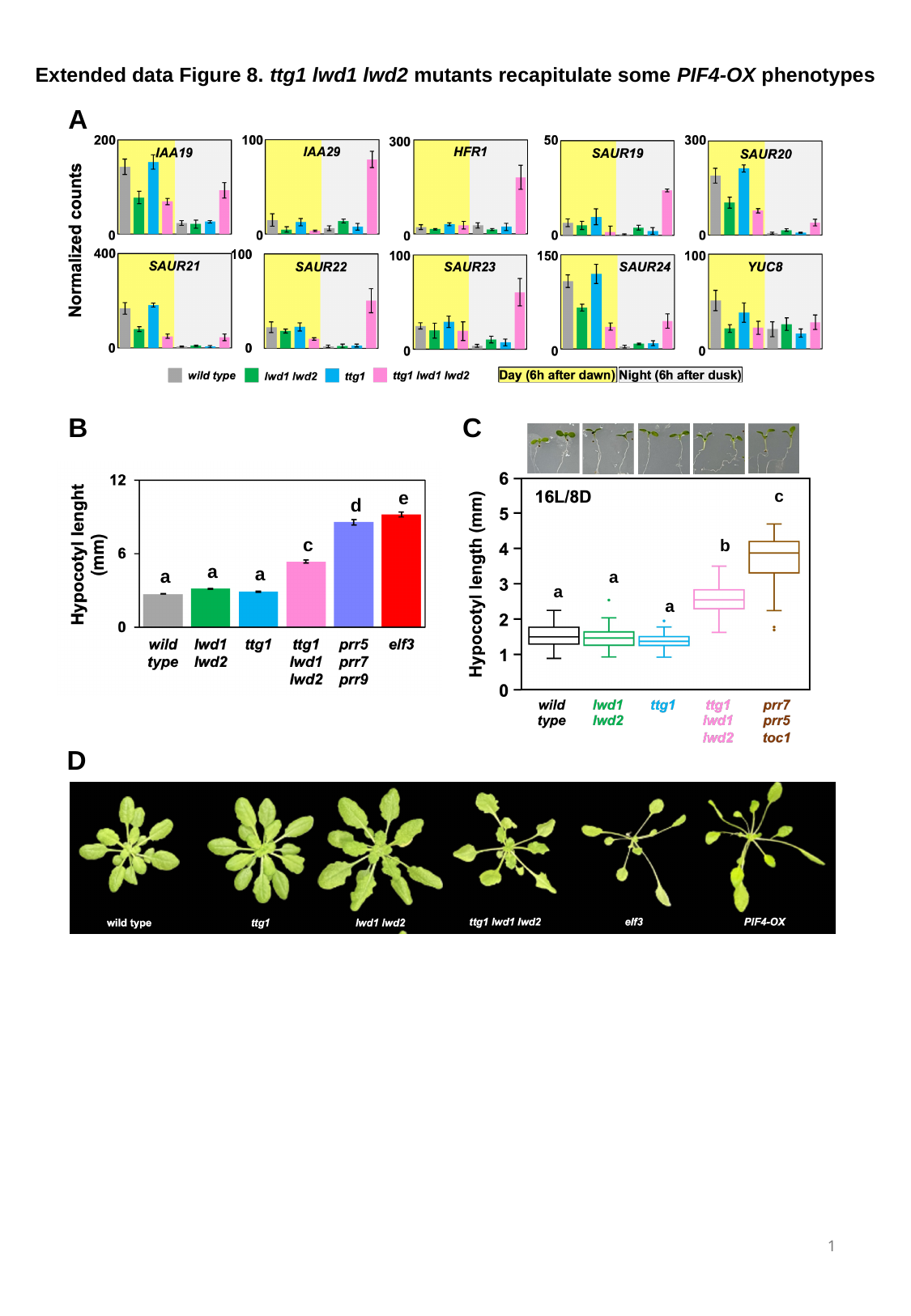

Extended data Figure 8. ttg1 lwd1 lwd2 mutants recapitulate some PIF4-OX phenotypes
A
B
C
c
b
a
a
a
e
d
c
a
a
a
D
 1
