## Supplementary Table S2 for "Arbor function of TRANSPARENT TESTA GLABRA 1 and LIGHT REGULATED WD scaffold proteins in the Arabidopsis circadian oscillators includes transcriptional repression through PSEUDO RESPONSE REGULATORS"

| **Line** | **Genetic Background** | **Source** |
| --- | --- | --- |
| wild type *pCCA1::LUC* | Col-0 | [27] |
| *ttg1* *pCCA1::LUC* | Col-0 | [27] |
| *lwd1 lwd2* *pCCA1::LUC* | Col-0 | [27] |
| *ttg1 lwd1 lwd2* *pCCA1::LUC* | Col-0 | [27] |
| *elf3 pCCA1::LUC* | Col-0 | This study |
| *prr5 prr7 toc1 pCCA1:UC* | Col-0 | NASC N2107712 |
| Wild type *pTOC1:LUC* | Col-0 | [52] |
| *ttg1 lwd1 lwd2* *pTOC1::LUC* | Col-0 | This study |
| wild type *pPRR7:LUC* | Col-0 | [52] |
| *ttg1 lwd1 lwd2* *pPRR7::LUC* | Col-0 | This study |
| *pCCA1::CCCA1-YFP* | Col-0 | [31] |
| *lwd1 lwd2* *pCCA1::LUC* pLWD1::LWD1-Venus::LWD1dss | Col-0 | This study |
| *ttg1 pCCA1::LUC pTTG1::Venus-TTG1::LWD1dss* | Col-0 | This study |
| *lwd1 lwd2* pLWD1::LWD1-LUC::LWD1dss | Col-0 | This study |
| *ttg1 pTTG1::TTG1-LUC::LWD1dss* | Col-0 | This study |
| Wild type *pTOC1::LUC* | Ws | NASC N9960 |
| *lhy cca1 pTOC1::LUC* | Ws | NASC N2107351 |
| *elf3 pCCA1::LUC* | Col-0 | This study |
| *PIF4-OX* | Col-0 | [59] |

**Table S2. *Arabidopsis thaliana* lines used in this study.**

**Source references in main text**
