## Supplementary Table S3 for "Arbor function of TRANSPARENT TESTA GLABRA 1 and LIGHT REGULATED WD scaffold proteins in the Arabidopsis circadian oscillators includes transcriptional repression through PSEUDO RESPONSE REGULATORS"

| Gene target | Forward primer | Reverse primer |
| --- | --- | --- |
| PP2A | TATCGGATGACGATTCTTCGTGCAG | GCTTGGTCGACTATCGGAATGAGAG |
| UBQ10 | GGCCTTGTATAATCCCTGATGAATAAG | AAAGAGATAACAGGAACGGAAACATAGT |
| PIF4 | GCGGCTTCGGCTCCGATGAT | AGTCGCGGCCTGCATGTGTG |
| PIF5 | GGGGTACAATCATCTCCATACAT | CCATGTACCTAGCGAGCTGCTCC |
| CDF5 | TGGTCTCCGAAACTTCTCAC | GCAACTTCATCACACAATGG |

**Table S3. qPCR primers**
